# Resolving the nanoscale structure of β-sheet assemblies using single-molecule orientation-localization microscopy

**DOI:** 10.1101/2023.09.13.557571

**Authors:** Weiyan Zhou, Conor L. O’Neill, Tianben Ding, Oumeng Zhang, Jai S. Rudra, Matthew D. Lew

## Abstract

Synthetic peptides that self-assemble into cross-β fibrils have remarkable utility as engineered biomaterials due to their modularity and biocompatibility, but their structural and morphological similarity to amyloid species has been a long-standing concern for their translation. Further, their polymorphs are difficult to characterize using spectroscopic and imaging techniques that rely on ensemble averaging to achieve high resolution. Here, we utilize single-molecule orientation-localization microscopy (SMOLM) to characterize fibrils formed by the designed amphipathic enantiomers, KFE8^L^ and KFE8^D^, and the pathological amyloid-beta peptide Aβ42. SMOLM reveals that the orientations of Nile red, as it transiently binds to both KFE8 and Aβ42, are consistent with a helical (bilayer) ribbon structure and convey the precise tilt of the fibrils’ inner and outer backbones. SMOLM also finds polymorphic branched and curved morphologies of KFE8 whose backbones exhibit much more heterogeneity than those of more typical straight fibrils. Thus, SMOLM is a powerful tool to interrogate the structural differences and polymorphism between engineered and pathological cross β-rich fibrils.

## Introduction

Peptides that self-assemble into cross-β fibrils are an exciting class of biomaterials due to the ease of designing their sequences and their ability to present functional ligands appended to their N- or C-terminus on the fibril surface^1^. In addition to their ability to assemble in aqueous physiological buffers, these structural scaffolds have powerful functionalities for applications in tissue engineering, regenerative medicine, drug delivery, bioorganic semiconductors, and biocatalysis^2^. In peptides with strictly alternating polar and non-polar residues, hydrophobic and hydrophilic residues are segregated on the opposite sides of the sheet, providing an amphiphilic surface that drives assembly into supramolecular cross-β fibrils^3^. However, many cross β-fibrillizing peptide sequences derived from pathological amyloid proteins do not strictly adhere to the alternating polar/non-polar design, and the role of cross-β fibrils in amyloid diseases is a topic of intense study and debate^4^. Thus, a wide range of natural or designed peptide sequences can adopt very similar or even the same cross-β structures, and once fibrillized, it can be very difficult to distinguish them based on their morphology^5^. Moreover, amphipathic designer peptides assemble into cross β-rich structures that closely resemble amyloids^6^. Therefore, understanding how toxic, benign, and functional fibrils differ in their molecular packing and nanoscale structure is paramount but not yet fully understood. Further, the polymorphic nature of amyloid fibrils adds another layer of complexity where a single peptide can form a range of molecularly distinct fibrils^7^.

Existing ultrahigh-resolution characterization tools, e.g., X-ray and electron diffraction methods, solid-state NMR spectroscopy, and cryo-EM^8–12^, can achieve atomic resolution bur require averaging over an ensemble. Either each polymorphic species must be isolated individually, or extensive datasets must be collected for computational processing—both of which are tremendously challenging. In addition, various sample preparation procedures (e.g., lyophilization and vitrification) can bias assemblies toward specific morphologies^9^. Both atomic force microscopy (AFM) and transmission electron microscopy (TEM) have been widely used for characterizing the morphologies and dimensions of fibrillar peptide assemblies but are blind to the molecular interactions and packing between peptides and the internal structure of self-assembled fibrils^13^.

In the last decade, single-molecule localization microscopy (SMLM) has been a powerful super-resolution fluorescence modality for quantitatively investigating the structural features of amyloids^13–16^. By repeatedly localizing “flashes” from individual fluorophores over time, SMLM has visualized the static and dynamic behavior of self-assembling fibrillar systems and their intermolecular interactions^17,18^. Beyond covalent labeling, binding-activated blinking^19^ of environment-sensitive amyloidophilic dyes can also be used for SMLM. Termed transient amyloid binding (TAB)^20^, a variety of dyes, including p-TFAA^21^, Nile red^22–24^, thioflavin T^20^, thioflavin X^25^, SYPRO orange, and LDS772^26^, can visualize self-assembling fibrillar systems under physiological conditions for hours to days. However, with localization precisions of ∼10 nm^20,22–25,27^, TAB SMLM cannot interrogate nanometer-scale details.

To overcome this limitation, combined imaging of fluorophore positions and orientations^28^, termed single-molecule orientation-localization microscopy (SMOLM), is actively being developed and reveals remarkable heterogeneity in fibrillar nanostructures with exquisite detail^22,23,26,29,30^. Orientation precisions of 4 ° -8 ° are possible when imaging dim emitters (∼500 photons)^23^, and when using bright fluorophores (∼5000 photons), 2.0° orientation precision in 3D can be achieved^29^. SMOLM’s high measurement sensitivity thus has great potential for visualizing the nanoscale details of various polymorphs of β-sheet fibril assemblies.

In this study, we utilized SMOLM to understand spatial organization of constituent peptides in cross-β fibrils generated from the designer amphipathic sequence KFE8 (Ac-FKFEFKFE-NH_2_) and Aβ42, the etiological agent of Alzheimer’s Disease. KFE8 assembles into fibrils with high solubility, and numerous spectroscopic studies have explored the impacts of sequence length and pattern variation on assembly. Moreover, initial assembly conditions exert a significant influence on the polymorphic nature of KFE8 fibrils and its potential biological interactions. Our orientation data reveal that the architectures of both KFE8 and Aβ42 are consistent with a helical bilayer ribbon, with KFE8 showing clear helicity in its measured backbone tilt angles and Aβ42 resembling an approximate straight line with modest backbone tilt; these structural features are hidden in standard SMLM. In addition, the rotational “wobble” of NR is significantly smaller and more consistent along the length of Aβ42 fibrils compared to those of KFE8, thereby demonstrating the exquisite sensitivity of TAB dyes for visualizing structural details of cross-β fibrils. SMOLM orientation data also show that KFE8 fibrils exhibit much more structural variability than those of Aβ42, despite the relative simplicity of the KFE8 sequence. Finally, SMOLM revealed rare branching, enclosed loop, and curved morphologies of KFE8, all of which were prepared under identical conditions to typical straight fibrils. However, these unusual morphologies displayed greater heterogeneity and structural diversity, as measured by NR orientations, thus demonstrating the power of SMOLM for uncovering and characterizing structural polymorphism.

## Results

### Visualizing β -sheet assemblies using transient binding-activated fluorescent probes

We performed SMOLM on homochiral KFE8^L^, KFE8^D^, and 42-residue amyloid-β peptide (Aβ42) assemblies using Nile red (NR, Fig. 1a, Inset) via the transient binding-activated blinking mechanism (see Materials and methods for details). Due to its solvatochromic and fluorogenic response^31,32^, NR remains dark in water but emits strong fluorescence upon binding to the β -sheet assemblies (Fig. 1a). Fluorescence “flashes” from NR coincide with binding and unbinding events, and an ample supply of NR molecules in solution enables a stable rate of blinking NR to be localized along each peptide assembly, whereby even hours-long SMOLM imaging is possible^20,33^. Here, Fig. S1 shows that the number of blinking SMs, i.e., localizations, is stable in a typical imaging experiment, which lasts for 100 s when we capture 5000 frames of each field of view (FOV).

**Fig. 1.**
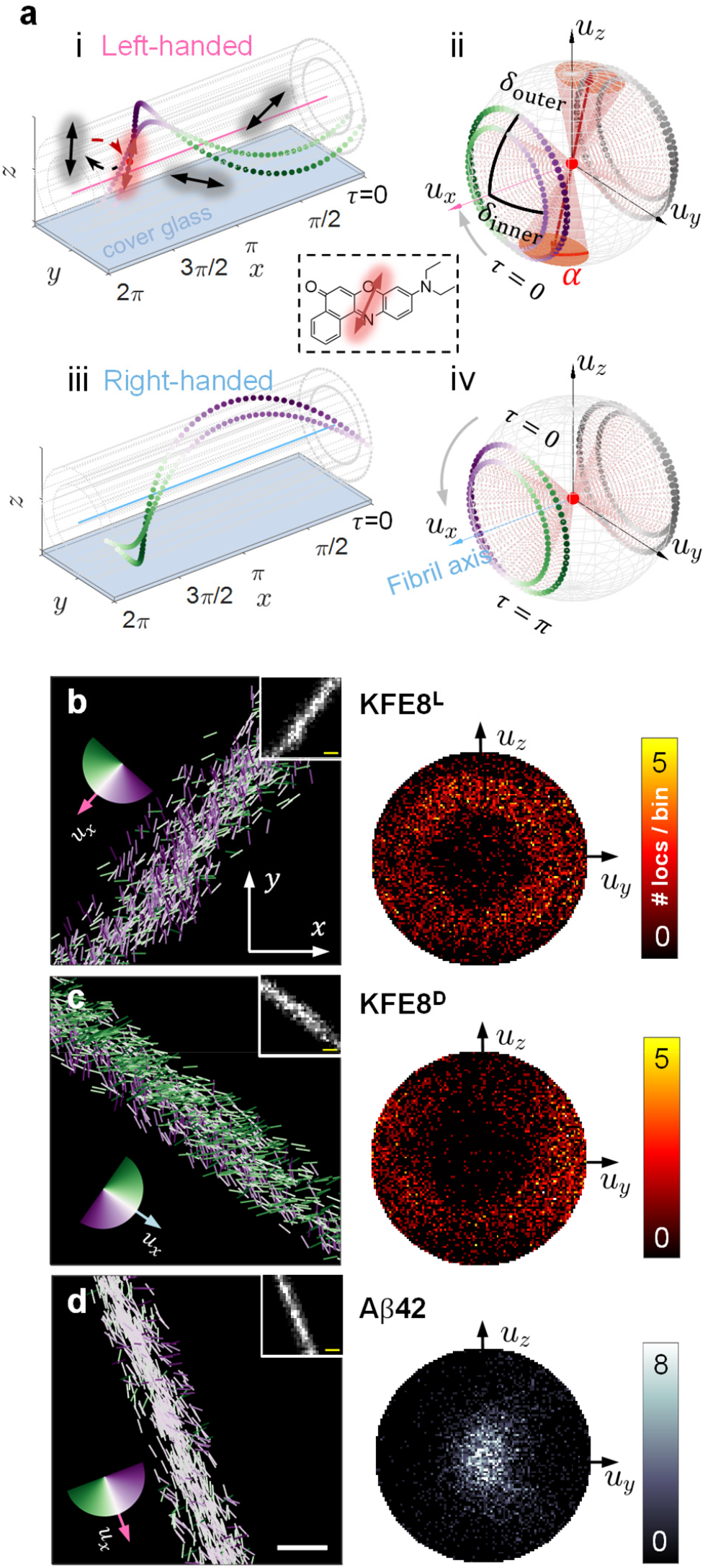
Single-molecule orientation-localization microscopy (SMOLM) of Nile red reveals the nanoscale structure of KFE8 and Aβ42 fibrils. **a**, (i) Transient binding of fluorogenic probes (red double-headed arrow) to a left-handed (LH) helical bilayer. Nile red (NR) binds to the helical bilayer parallel to its backbone, which exhibits a periodic helical twist along its long axis. NR is nonfluorescent in solution (dark arrows). (*x, y, z*), spatial coordinates defined by the microscope. *τ*, helix phase, which for a helical structure, also corresponds to the height of the fibril above the coverglass. *Inset*, chemical structure of NR. (ii) A model orientation distribution of NR accumulated across all possible binding sites along the helical bilayer. (*u*_*x*_, *u*_*y*_, *u*_*z*_), orientation coordinates defined by the long axis *u*_*x*_ of the fibril. *α*, wobble cone half angle of NR, quantifying its orientation freedom during a camera frame. *δ*_inner_ and *δ*_outer_, inner and outer backbone tilt angles of the helical bilayer, respectively. *τ* = 0 corresponds to a minimum value of *u*_*z*_ and increases in the direction of the gray arrow. (iii and iv) Right-handed (RH) helical bilayer and the corresponding orientation distribution of NR. *τ* = 0 corresponds to a maximum value of *u*_*z*_ and increases in the direction of the gray arrow. **b-d**, (Left) Experimental SMOLM images and (Right) their corresponding 3D orientation distributions on the *u*_*x*_ ≥ 0 hemishpere for (**b**) KFE8^L^ (LH), (**c**) KFE8^D^ (RH)^34^, and (**d**) Aβ42 (LH)^37,48^. Each NR localization is depicted as a line segment whose color and orientation are set by the projection of the molecule’s 3D orientation into the *xy* plane relative to the long axis *u*_*x*_ of the fibril. *Insets*, superresolution SMLM images. Scale bars are 100 nm. The orientation distributions are binned using *u*_*y*_ = (-1,-0.98,…,1) and *u*_*z*_ = (-1,-0.98,…,1).

Without orientation information, standard single-molecule localization microscopy produces nearly indistinguishable images of KFE8^L^, KFE8^D^, and Aβ42 (Fig. 1b-d, Inset and Fig. S2a-c, Inset), though KFE8^L^ and KFE8^D^ fibrils are wider than Aβ42 (Fig. S2d-e, full width of the half maximum (FWHM) of 60 – 110 nm for KFE8^L^ and KFE8^D^ and 30 – 70 nm for Aβ42). According to AFM and TEM measurements, KFE8^L^ and KFE8^D^ have a width of ∼8 nm and a pitch of ∼20 nm^34–36^, while the width and the pitch of Aβ42 are ∼9 nm and ∼100 nm, respectively^37–39^. Thus, it is almost impossible to reveal the putative ribbon structure and quantify the polymorphs using the conventional SMLM.

However, measuring the location and orientation of each blinking fluorophore simultaneously has great potential for revealing supramolecular structural details. Recent SMOLM studies have shown that amyloidophilic dyes like NR bind to β-sheet fibrils with their transition dipoles parallel to the fibril backbone (Fig. 1a i)^22,23,40,41^, which is consistent with detailed simulation studies of their binding modes^42–44^. If NR molecules bind to grooves along a helical fibril, they will be tilted with respect to the long axis of that fibril (Fig. 1a i). We denote this angle as the backbone tilt angle *δ* (Fig. 1a ii); larger diameter-to-pitch ratios lead to larger backbone tilt angles. Thus, accumulated over the length of the fibril, the orientations of NR measured via SMOLM will form a circle on the orientation sphere, where the *u*_*x*_ direction is defined to be parallel to the fibril’s long axis. Due to dipole symmetry (Fig. 1a ii), the *u*_*x*_ < 0 hemisphere is a replica of the *u*_*x*_ ≥ 0 hemisphere. Therefore, only the *u*_*x*_ ≥ 0 hemisphere is considered in the following sections.

In this work, SMOLM resolves remarkable differences between these β -sheet-formed fibrils, where the location and the orientation of each NR molecule are depicted as the center and the direction, respectively, of each line in Fig. 1b-d. The lines are color-coded according to the relative angle between the orientation of NR (i.e., its 3D orientation projected into the *xy* plane) versus the long axis *u*_*x*_ of the fibril. These data are collected by a custom-built polarization-sensitive microscope using a vortex phase mask^23^ and analyzed by a bespoke regularized maximum likelihood estimator, RoSE-O^45,46^ (see Materials and Methods for details).

SMOLM images show that NR molecules bound to KFE8^L^ and KFE8^D^ span a large range of orientations (Fig. 1b and c, Left), while NR orientations are concentrated near the white region (Fig. 1d, Left), i.e., parallel to the fibril axis *u*_*x*_, for Aβ42. The distinct structural characteristics of KFE8 and A β 42 are clearly revealed in their corresponding orientation distributions (Fig. 1b-d, Right). Both KFE8 systems have a bright ring-like distribution with a dark center (Fig. 1b and c, Right). Conversely, Aβ42 has a bright center with a dark outer boundary (Fig. 1d, Right). Given the connection between NR orientations and the tilt angle *δ* of the fibril’s helical backbone, these data clearly show that the KFE8 fibrils in Fig. 1*b* and *C* have larger tilt angles *δ* than Aβ42 does. Therefore, measuring orientations using SMOLM sheds new light on the nanoscale details of supramolecular structures and possible polymorphism^14,15,47^ beyond what SMLM is capable of. We next quantify how the observed circular orientation distributions are related to the helicity of a supramolecular structure and the number of layers it contains.

### Experimental characterization of helical ribbon supramolecular structures using SMOLM

The left-handed helical supramolecular structure of KFE8^L^ was first observed using AFM and TEM over 20 years ago^35^, and molecular dynamics simulations suggest that the complementary amphipathic β - strands of KFE8 form a bilayer, i.e., a helical ribbon^35,49^. Since then, the putative helical ribbon model has been widely accepted^36,50–52^. To date, no further direct experimental validation has been realized.

Fig. 2a shows the SMOLM orientation distributions of KFE8^L^, KFE8^D^, and A β 42 accumulated from 44 (KFE8^L^), 38 (KFE8^D^), and 32 (Aβ42) isolated linear fibrils (example fibrils are shown in Fig. 1b-d and Fig. S2a-c). Similar to the individual fibrils in Fig. 1b-d, we find a bright ring-like distribution with a dim center for KFE8^L^ (Fig. 2a, Left) and KFE8^D^ (Fig. 2a, Middle) and a bright center and a dim boundary in Aβ42 (Fig. 2a, Right).

**Fig. 2.**
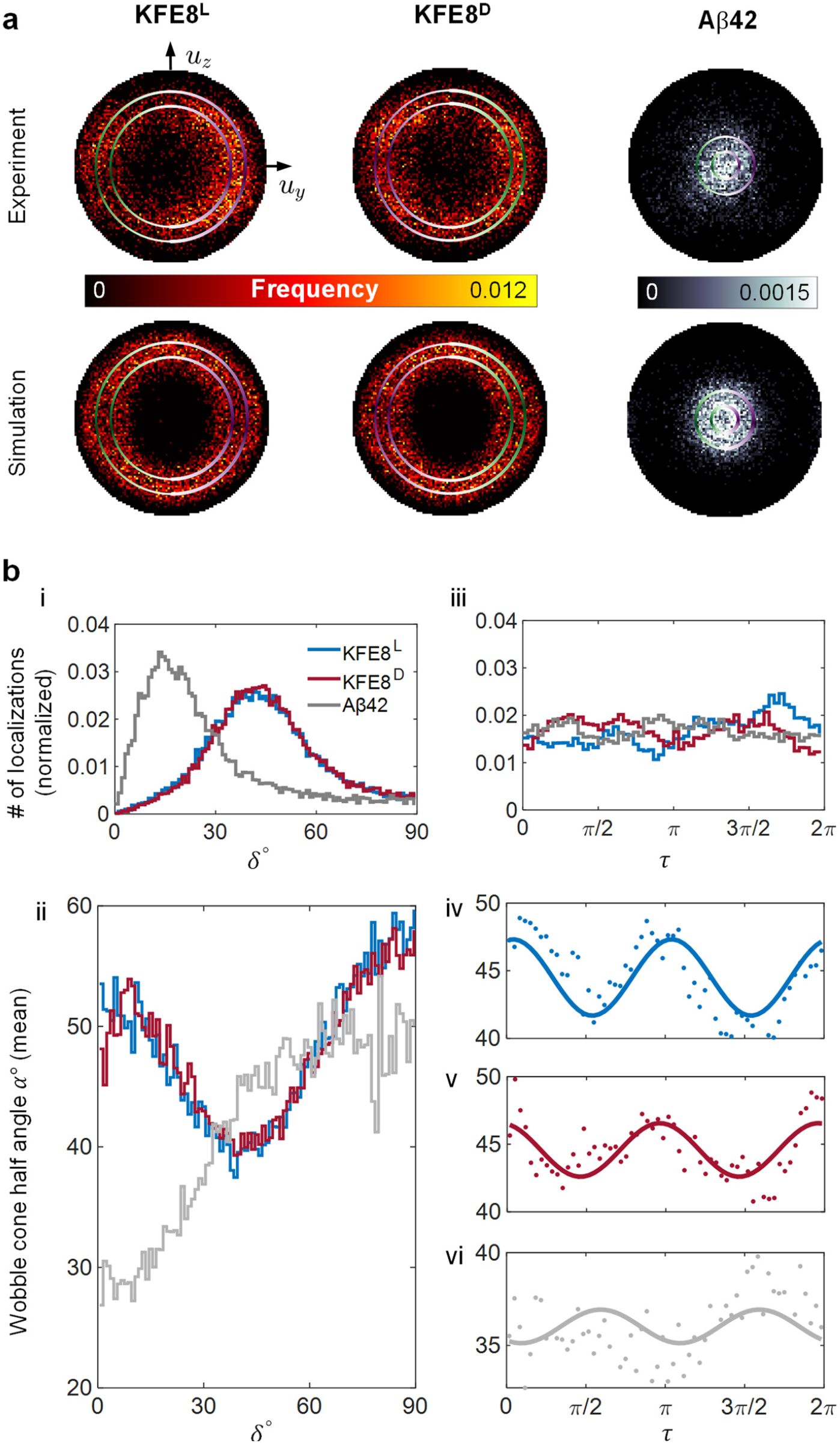
NR orientation distributions, localization densities, and wobble shed light on the helical bilayer structure of KFE8 and Aβ42 fibrils. **a**, (Top) Experimental and (Bottom) simulated orientation distributions of (Left) KFE8^L^, (Middle) KFE8^D^, and (Right) Aβ42 with overlaid backbone tilt angles *δ*_inner_ and *δ*_outer_ calculated using our helix optimization algorithm (See SI Fig. 4). The circles, i.e., the measured backbone tilt angles, are color-coded with respect to helix phase *τ* (See Fig. 1a). The orientation distributions are binned using *u*_*y*_ = (-1,-0.98,…,1) and *u*_*z*_ = (-1,-0.98,…,1). Filter: signal ≥ 500 photons, wobble cone half angle *α* ≤ 30°. **b**, NR (i, iii) localization density and (ii, iv-vi) wobble quantified as a half-cone angle *α* in degrees. Filter: signal 500 photons. (i and ii) Localization density and wobble are quantified with respect to backbone tilt angle *δ*. Bin size: 1°. (iii-vi) Localization density and wobble are quantified with respect to helix phase *τ*. (iv-vi) SMOLM measurements are represented as points; solid lines are from fits to sinusoidal functions. Bin size: *π*/30 (6°). Data are accumulated from all measured fibrils. See Figs. S8 and S9 for full distributions.

To determine supramolecular structure from SMOLM data, we devised a helix optimization algorithm (Materials and methods), which translates SMOLM orientation measurements into tilt angle estimates (Fig. S4a iii and iv). Depending on the choice of model, the algorithm finds either 1) a pair of tilt angles (*δ*_inner_, *δ*_outer_) of a helical bilayer (Eq. 5), or 2) a single tilt angle *δ*_mono_ (Eq. 6) that best “fits” the NR orientations observed via SMOLM. To validate model selection and performance of the optimization algorithm, we simulated the process of imaging NR with specific orientation distributions from helices (Fig. S4a i) using the polarized vortex SMOLM (Fig. S4a ii). Briefly, in the case of the helical bilayer model, we treat the pair of measured backbone tilt angles (Fig. S4a iv) as ground truth. Then, we uniformly sample two circles on the orientation hemisphere that correspond to NR molecules bound to the double-layer helical ribbon. Using vectorial diffraction theory^53^, we simulate images formed by the polarized vortex microscope that are corrupted by Poisson shot noise, and from these images, NR orientations are estimated by RoSE-O^45^. The resulting orientation distributions, accumulated over all NR orientations spanning both layers of a helical ribbon, are a realistic representation of how the polarized vortex SMOLM would perform when imaging a helical ribbon with known backbone tilt angles.

We performed extensive simulations to validate the robustness of our helix optimization strategy. First, we simulated 20 helical ribbons with similar backbone tilt angles as KFE8 and found that the algorithm has high accuracy and precision (Table S1, 5040 SMs per ribbon matching our experiments). The bias in measuring backbone tilt is less than 1° and 1.5° in 87.5% and 97.5% of the cases, respectively, while our precision in estimating both *δ*_inner_ and *δ*_outer_ is less than 1.1°. Second, we found that the algorithm is robust against the number of SMs (Fig. S6a). The standard deviation is less than 0.5° and the bias is less than 0.65° even when the number of SMs is only around 300. When the number of SMs reaches 5000, the standard deviation reduces to 0.2° and the bias is 0.59 °. Third, to investigate the generality of SMOLM’s ability to measure helical structures, we simulated monolayer helices with backbone tilt angles *δ* ranging from 0° to 90° (Fig. S6b). When *δ* ∈ [ 10 °, 80 ° ], the helix optimization algorithm shows a bias less than 1.1°, with precision less than 0.1° (630 SMs per helix, 10 helices with same size for precision quantification). For helices with 20 nm pitch, *δ* = 10° corresponds to a radius of 0.56 nm and *δ* = 80° to 18.05 nm. We found that the bias is monotonic outside this range and can be corrected using these data to restore accuracy. Due to computational cost, we did not quantify the performance of measuring *δ*_inner_ and *δ*_outer_ for helical ribbons across the entire range, 0° to 90°. Overall, our simulations of SMOLM and its associated orientation analysis show excellent performance for helical structure determination.

Using our algorithm, we found that the backbone tilt angles of the inner and the outer layer of KFE8^L^ are *δ*_inner_ = 38.90° and *δ*_outer_ = 50.34° (Fig. 2a, Top left, 5194 total SMs). Assuming a pitch of 20 nm^34–36^, the corresponding radii of the inner and outer layer are 2.57 nm and 3.84 nm, respectively. The backbone tilt angles of KFE8^D^ are *δ*_inner_ = 39.20° and *δ*_outer_ = 50.58° (Fig. 2a, Top middle, 5104 total SMs), corresponding to inner and outer layer radii of 2.60 nm and 3.87 nm, respectively. These measurements are consistent with TEM^34^. In contrast, the tilt angles of the inner and the outer layer of Aβ42 are *δ*_inner_ = 7.86° and *δ*_outer_ = 16.96° (Fig. 2a, Top right, 4646 total SMs). Remarkably, the experimental NR orientation distributions (Fig. 2a, Top) match our simulated orientation distributions of a helical ribbon (Fig. 2a, Bottom) for both KFE8 and Aβ42. On the other hand, simulated orientation distributions of a helical monolayer (Fig. S5c) are much narrower than the experimental distribution even when accounting for noise stemming from photon counting and a finite number of NR localizations. Therefore, SMOLM shows that NR binds to these β-sheet fibrils in a manner consistent with double-layer ribbons instead of single-layer tapes.

### Structural insights of β -sheet assemblies revealed by fluorogenic probes

Since NR molecules emit fluorescence only when bound to each fibril, we posited that the photophysical and rotational behavior of each NR molecule could reveal insights into the structure of each β-sheet assembly. Thus, we quantified the localization density and the “wobble” cone half angle *α* (Fig. 1a, ii) of the SMs as a function of backbone tilt angle *δ*. We note that our helical model features a one-to-one mapping between backbone tilt angle (Eqs. 2 and 4) and molecule position (Eqs. 1 and 3) relative to the fibril’s central axis. Thus, for a given fibril, a NR molecule with relatively small *δ* may be interpreted as binding to an “interior” binding site close to the central axis, while large *δ* correspond to “exterior” sites that are relatively far from the axis (Fig. 1a, pink and blue axes).

Enantiomeric KFE8^L^ and KFE8^D^ feature almost identical localization density curves (Fig. 2b i, KFE8^L^: 24715 total localizations, KFE8^D^: 25283 total localizations, Aβ42: 15708 total localizations), which means that NR binds similarly to KFE8^L^ and KFE8^D^ even though they have opposite handedness. We find that 38% and 37% of the SMs bound to the interior of KFE8^L^ ( *δ*≤38.90° ) and KFE8^D^ ( *δ*≤ 39.20°), respectively, while 29% and 30% of the SMs visited the region between the two layers of KFE8^L^ ( 38.90° < *δ* < 50.34° ) and KFE8^D^ ( 39.20° < *δ* < 50.58°), respectively. The rest (33% for both KFE8^L^ and KFE8^D^) exhibited tilt angles consistent with the exterior of the ribbon (KFE8^L^: *δ* ≥ 50.34° and KFE8^D^: *δ* ≥ 50.58°). In contrast, NR gives a distinctly different localization density distribution when binding to Aβ42 fibrils. We observed 10%, 27%, and 63% of the localizations to be within the inner layer ( *δ*≤ 7.86°), between the bilayer (7.86° < *δ* < 16.96°), and from the outer layer (*δ* ≥ 16.96°), respectively. The peak localization densities correspond to backbone tilt angles of 41° for KFE8^L^, 44° for KFE8^D^, and 13° for Aβ42. The larger angles *δ* for homochiral KFE8 imply that they have a larger diameter-to-pitch ratio than Aβ42, which is consistent with previous studies of KFE8^34,35,49,51^ and Aβ42^54,39,37,38^.

We next examine SMOLM measurements of NR “wobble” while bound to each fibril, quantified as a half-cone angle *α* explored by NR during a camera frame (Fig. 1a, ii). We find remarkable similarity between the enantiomeric KFE8 pairs (Fig. 2b, ii), suggesting that opposite handedness does not significantly affect the local binding behavior of NR to each assembly. NR shows a minimum wobble when *δ* ∈ [38°, 50°], which corresponds to the region between the two layers of the helical ribbon. Notably, this trend is in good agreement with the helical ribbon model of KFE8, where the phenylalanine residues are buried inside to form a hydrophobic core. NR exhibits stronger and more frequent binding in this hydrophobic core versus the more polar exterior, and SMOLM consequently detects a greater number of NR molecules within this range of tilt angles with a commensurate minimum in rotational freedom. In stark contrast for Aβ42 (Fig. 2b, ii), NR demonstrates systematically larger wobble for larger tilt angles, which correspond to binding to the helix exterior. When *δ*≤16.96°, i.e., inside the inner layer and between the bilayer, the wobble angle of NR is at a nearly stable minimum near 30° . This small rotational freedom implies that the side-chain grooves of Aβ42 fibrils more strongly constrain the rotation of NR than those of KFE8 systems, thereby demonstrating the potential of SMOLM for sensing the structure and chemical environments near β-sheet assemblies.

Interestingly, for both KFE8 and Aβ42, we find that the localization density of NR reaches a maximum (Fig. 2b, i) at nearly the same backbone tilt angles as where the wobble of NR attains its minimum (Fig. 2b, ii). Specifically, NR with orientations further from the most frequently measured backbone tilt angles (*δ* ∈ [38°, 50°] for KFE8 and *δ* ∈ [8°, 17°] for Aβ42) exhibit larger wobble than those whose tilt angles lie near the peak. We interpret this correlation to be consistent with the TAB blinking mechanism of NR (Fig. 1a); when NR molecules are further away from the assemblies, they have more rotational freedom. In addition, the surrounding chemical environment is more polar, leading to a smaller quantum yield. Thus, for β-sheet assemblies, smaller NR wobble is likely directly linked to more hydrophobic binding sites, which in turn yield more frequent NR localizations.

We next characterize β-sheet assemblies along the helical trajectory, parameterized by phase *τ* (with *τ* ∈ [0, 2*π*) defined as in Fig. 1a, i-iv), by quantifying NR localization density and rotational freedom. Interestingly, we find that NR binds with near uniform frequency along each fibril, regardless of the variant (Fig. 2b, iii). Surprisingly however, we find striking sinusoidal variations in the half cone angle *α* as a function of helix phase *τ* (Fig. 2b, iv-vi), i.e., as the fibrils rise and fall with respect to the coverslip (Fig. 1a, ii and iv). Fitting the wobble angle measurements to sinusoid functions yields *α*_L_ = 2.8° cos(2*τ*-0.25)+ 44.5° for KFE8^L^, *α*_D_ = 2.0° cos(2*τ*+ 0.24)+ 44.6° for KFE8^D^, and *α*_A42_ = -0.9° cos(2*τ*-0.58)+ 36.0°; we constrain the frequency of each sinusoid to ensure continuity over *τ* . The magnitude of these variations for KFE8 is more than twice as large as those observed in simulated imaging experiments where *α* is held constant (Fig. S7a).

Notice that the wobble angle *α* oscillates twice over a single helical period for KFE8^L^ and KFE8^D^ (Fig. 2b, iv-vi), achieving minima near *τ* = *π*/2 and *τ* = 3*π*/2 (Fig. 2b, iv and v); this minimum wobble occurs where the helices are either in contact with the glass coverslip or are furthest away from the coverslip. In addition, we observe that the average wobble of NR is almost the same when binding to KFE8^L^ and KFE8^D^, but the variations in wobble along the helix are 40% larger for KFE8^L^ than for KFE8^D^. In contrast, NR exhibits different wobble behavior when bound to Aβ42 (Fig. 2b, vi). As opposed to reaching minima as in KFE8, NR rotational freedom is maximized for sections of Aβ42 fibrils that are close to and far away from the coverglass, albeit with variations that are approximately half as large for Aβ 42 compared to KFE8. One possible reason for this difference is Aβ42’s smaller backbone tilt angles (Fig 2a, Top), which means that its fibrils are more approximate to straight lines instead of helices. Thus, helix-related variations are simply less significant for Aβ42 fibrils.

### Nile red orientations reveal polymorphism across fibrils of KFE8 and Aβ42

Characterizing polymorphism of self-assembling peptides is critical to the rational design of biomaterials^5,55^ and effective drug-delivery vehicles^56,57^. Polymorphism has been studied in both synthetic and natural systems^58–60^, but structure-function relationships remain difficult to characterize due to methodological challenges. Motivated to unmask possible polymorphism, we quantified backbone tilt angles *δ* and wobble cone half angles *α* of NR bound to linear and isolated fibrils of KFE8^L^, KFE8^D^, and Aβ42 across 90 fields of view (FOVs). Representative SMOLM images are shown in Fig. 1b-d and Fig. S2a-c. Sorting the FOVs according to the median backbone tilt angle, we observe remarkable variations in measured backbone tilts of KFE8 (Fig. 3a and b) that are hidden when averaging data across all fibrils (Fig. 2). The medians of the backbone tilt angle *δ* within each FOV ranges from 35° to 48° for KFE8^L^ (Fig. 3a), 38° to 49° for KFE8^D^ (Fig. 3b), and 15° to 22° for Aβ42 (Fig. 3c). Surprisingly, despite its short sequence, fibrils of KFE8 display much more structural variability than those of Aβ42 in our experiments.

**Fig. 3.**
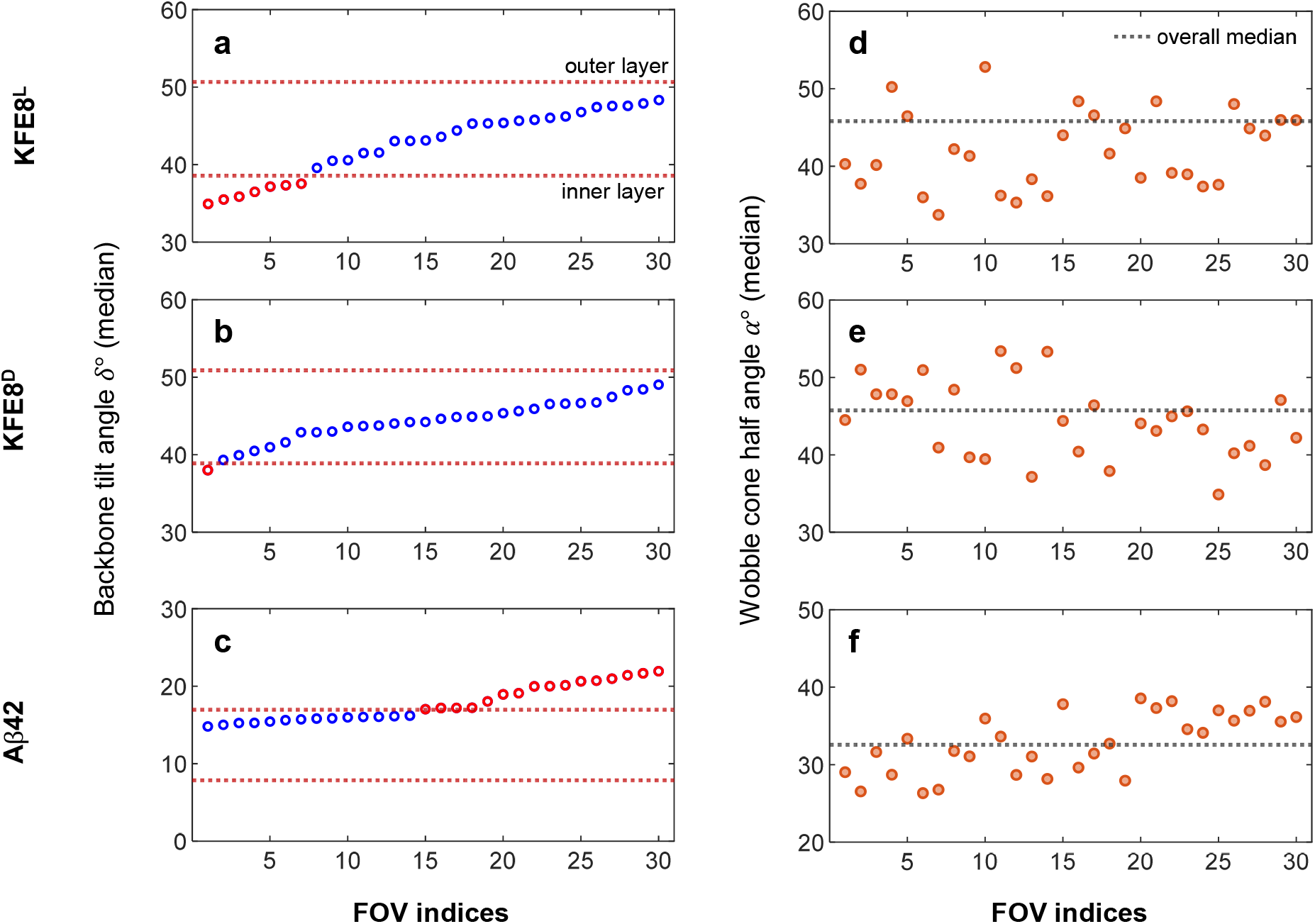
NR SMOLM quantifies heterogeneity among KFE8^L^, KFE8^D^, and Aβ42 fibrils. **a-c**, Median backbone tilt angles *δ* of NR measured within 30 fields of view (FOVs) for (**a**) KFE8^L^, (**b**) KFE8^D^, and (**c**) Aβ42. Hollow circles are median angles within each FOV. Red dotted lines are the backbone tilt angles *δ*_inner_ and *δ*_outer_ of KFE8^L^, KFE8^D^, and Aβ42, as estimated by helical optimization of data accumulated across all FOVs shown in Fig. 2a. **d-f**, Median wobble cone half angles *α* measured within 30 FOVs for (**d**) KFE8^L^, (**e**) KFE8^D^, and (**f**) Aβ42. Orange circles are the median angles within each FOV. Black dotted lines are median angles over all KFE8^L^, KFE8^D^, and Aβ42 FOVs in Fig. 2b. KFE8^L^ FOV 26 corresponds to fibril shown in Fig. 1b, FOV 4 to Fig. S2a, i, FOV 16 to Fig. S2a, ii, FOV 12 to Fig. S2a, iii, and FOV 21 to Fig. S2a, iv. KFE8^D^ FOV 30 corresponds to fibril shown in Fig. 1c, FOV 22 to Fig. S2b, i, FOV 15 to Fig. S2b, ii, FOV 18 to Fig. S2b, iii, and FOV 7 to Fig. S2b, iv. Aβ42 FOV 7 corresponds to fibril shown in Fig. 1d, FOV 29 to Fig. S2c, i, FOV 22 to Fig. S2c, ii, FOV 19 to Fig. S2c, iii, and FOV 10 to Fig. S2c, iv.

We also observe a large variation in NR wobble across FOVs (Fig. 3d-f; for full distribution, see Fig. S10). The overall median wobble is 46° for both KFE8^L^ and KFE8^D^ (Figs. 3d and e). However, NR wobble on each fibril tends to be more constrained, with 70% of the KFE8^L^ FOVs (Fig. 3d) and 63.33% of the KFE8^D^ FOVs (Fig. 3e) having a median smaller than that of the accumulated localizations. The analogous statistics for Aβ42 are more symmetric, with an overall median of 33° (Fig. 3f) and 53.33% of the FOVs having a median larger than that of the accumulated FOVs (Fig. 2b, vi).

Under physiological conditions, we observed not only linear fibrils but also rare KFE8 morphologies, including branching (Fig. 4a) and enclosed (Fig. 4d) structures (See Fig. S12 for more examples). Recently, branching has also been observed using high-speed atomic force microscopy^50^, but to our knowledge, enclosed, looping structures have never been reported. SMOLM images (Fig. 4a and d) depict NR orientations projected into the *xy* plane, which show striking diversity when bound to the underlying supramolecular helices. These detailed variations in structure are impossible to detect using other techniques.

**Fig. 4.**
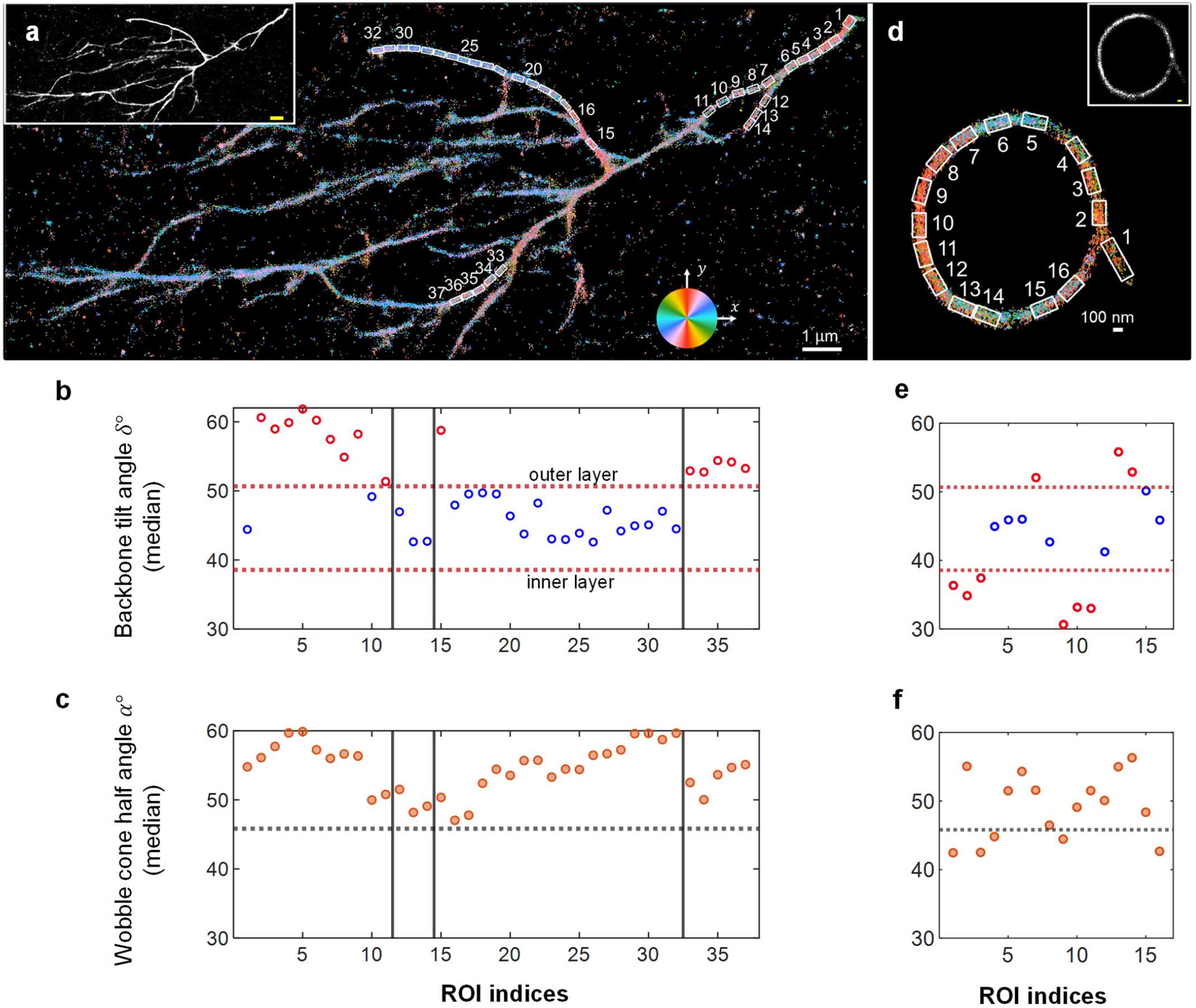
Structural polymorphism of KFE8^L^ revealed by SMOLM: **a-c**, Branching and **d-f**, enclosed loop structures. In **a-c**, two “parent” branches (ROIs 1-6 and 15-32) and three “child” branches (ROIs 7-11, 12-14, and 33-37) are chosen. (**a** and **d**) SMOLM and (Inset) SMLM images. NR orientations in the SMOLM images are represented as colored disks in **a** (108038 localizations [59% of the data] are randomly selected for plotting in **a** to avoid oversaturation) and colored lines in **d** (all data are shown). Each disk/line is color-coded according to the projection of the 3D NR orientation into the *xy* plane relative to the *μ*_*x*_ axis, which is defined by the microscope instead of the long axis of the fibril. (**b** and **e**) Median backbone tilt angles *δ* within each region of interest (ROI), which are shown in SMOLM images. Red dotted lines are the backbone tilt angles of the helical bilayer, as estimated by helical optimization of data accumulated across all FOVs shown in Fig. 2a, Left. Blue circles lie within the estimated backbone tilt angles of the bilayer; red circles outside the range. (**c** and **f**) Median wobble angle *α* (orange circles) within each ROI. Black dotted lines are the median of the accumulated experimental data in Fig. 2b. For more examples, see Fig. S12. Scale bars: (**a** and inset) 1 µm, (**d** and inset) 100 nm.

To investigate these polymorphs more closely, we analyzed individual segments along each structure, each with a length of 250 nm except ROI 1 in Fig. 4d, which was lengthened to 400 nm to ensure sufficient localization statistics. We found notable systematic deviations from the behavior of typical fibrils. First, the *median* backbone tilt angles *δ* of one parent branch (ROIs 1-11, Fig. 4b) are around 6° larger than the tilt *δ*_outer_ = 50.34° of the *outer layer* of typical KFE8^L^ fibrils (Fig. 3a). However, this trend does not hold for an offshoot child branch (ROIs 12-14). Another parent branch (ROIs 15-32) is comparable to typical assemblies, with median *δ*s that are between *δ*_inner_ and *δ*_outer_ . Second, we found that all measured NR wobble angles for the branching structure (Fig. 4c) are larger than those of linear fibrils. Thus, this branching KFE8 morphology contains binding sites that allow NR to rotate more freely, even while bound, than typical linear fibrils do.

We also find that the enclosed ring morphology of KFE8^L^ fluctuates along its circumference (Fig. 4e). Of the 16 regions we analyzed, 6 of them show median backbone tilt angles that are smaller than the tilt *δ*_inner_ of the *inner layer* of typical linear fibrils (Fig. 3a). These data imply that locally, these fibril segments have smaller diameters or longer periodicity than typical ones. Interestingly, three ROIs have larger tilt angles *δ* than *every* straight fibril we observed (Fig. 3a). We also observed varying degrees of NR rotational wobble along the entire structure (Fig. 4f), with 69% of the segments exhibiting a median *α* larger than that of typical linear fibrils. Analyses of similar enclosed and curved fibrils of KFE8^D^ reveal similar behaviors (Fig. S12); local segments show heterogeneous backbone tilts *δ* and rotational wobble *α*, some of which deviate from typical observations (Fig. 3b). Overall, SMOLM analysis shows that the morphologies of KFE8 assemblies exhibit strong variations, both between and within individual fibrils. Moreover, unusual morphologies, such as branching, looped, and curved structures show more heterogeneity than linear segments do.

## Discussion

Peptides that self-assemble into cross β-sheet fibrils have opened new perspectives in seemingly unrelated fields^2,61^. To gain molecular and structural insights into misfolded states of proteins involved in neurodegenerative diseases, scientists developed minimal peptide fragments capable of emulating the behavior of the parent protein. Peptides from natural proteins such as islet amyloid polypeptide (FGAILS)^62^, human calcitonin (DNFKFA)^63^, sup35 (GNNQQNY)^64^, and Aβ42 (KLVFFA)^65^ not only reveal structural information of the original systems, but also demonstrate that small peptides can self-assemble into supramolecular structures. They also have provided a deeper understanding of the formation of oligomeric intermediates and disease pathogenesis. Independently, a second family of designer peptides comprised of amphipathic sequences with strictly alternating hydrophobic and charged residues were first identified in *S. cerevisiae*^66^. This class of peptides also assemble into cross β-sheet fibrils and have attracted considerable interest as biomaterial scaffolds due to their ability to form hydrogels^67^. The close resemblance of these various cross β-rich structures directly contrasts with their disparate sequences and differing toxicity and function; the connections between molecular packing, structure, and biological functions are not yet fully understood.

The polymorphic nature of amyloid fibrils adds another layer of complexity, where a single peptide can form a range of molecularly distinct fibrils^7,60,68,69^. Structural polymorphs consistently manifest themselves as subtle alterations in the morphology and structure of β-sheet assemblies, arising from differences in inter or intra-residue interactions, the number of amino acid residues, their packing, and their orientations^69^. Computational and spectroscopic studies confirm that structural variations in pathological amyloid fibrils can be responsible for the observed disease variations in multiple neuropathies^70^. Interestingly, recent cryo-electron microscopy structural analysis of amphipathic designer peptides assembled under different sets of conditions identified the presence of multiple polymorphic species^36^. Whereas sequence variation of amphipathic peptides has been exploited to control long-range and bulk material properties such as crystallinity, gelation, morphology, and mechanical properties, reliable prediction of supramolecular structure from sequence information remains a significant challenge. Characterizing polymorphism is thus critical to elucidating structure-function relationships as well as accelerating the rational design of functional biomaterials.

Conventional bulk analysis and ensemble-based measurements may mask structural heterogeneity between and within individual fibrils and therefore be blind to rare species. In this work, we use NR TAB to non-covalently label KFE8 and Aβ42, thereby enabling SMOLM to characterize individual β-sheet fibrils with single-molecule sensitivity. NR orientation distributions accumulated across a multitude of isolated straight fibrils are consistent with a helical bilayer ribbon model (Fig. 2), and a bespoke helix optimization algorithm measures the associated backbone tilt angles *δ*_inner_ and *δ*_outer_ with excellent precision and accuracy (Figs. S6, S7, and Table S1). The larger degree of polymorphism in KFE8 versus Aβ42 is obvious when comparing the median backbone tilt angle and wobble of NR in each field of view individually. This remarkable variation (Fig. 3) is otherwise hidden by ensemble-averaged measurements (Fig. 2). Finally, our analysis of branching, enclosed looping, and curved morphologies of KFE8 shows that their fibrils are distinct (Figs. 4 and S12) from those of straight fibrils, thereby providing a first link between nanoscale helical ribbons and micron-scale morphology. Almost universally, we found significant variations both within and between individual fibrils, thereby underscoring the necessity of characterizing these structures in aqueous physiological buffers with the single-fibril sensitivity and nanoscale resolution afforded by TAB SMOLM.

We note that the structural details in our study are powerfully and uniquely conveyed by the connection between a supramolecular structure and the orientations of its binding sites for amyloidophilic probes (Fig. 1a). While the binding orientations of NR to Aβ42 fibrils are consistent with a double-layer ribbon model (Fig. S5c), we can also interpret these data as two types of binding sites that are linked to two different backbone tilt angles. Since high-resolution ssNMR and cryo-EM show S-shaped conformations formed by the stacking of Aβ42 peptides^9^, we hypothesize that the backbone tilts *δ*_inner_ and *δ*_outer_ measured by SMOLM correspond to two helical grooves within Aβ42 fibrils. Further theoretical and experimental investigations are necessary to determine the precise binding behaviors of NR.

As such, it is important to expand the library of TAB-compatible fluorophores^25,26^ to both probe new dye-structure interactions and to broaden the applicability of TAB SMOLM to new classes of peptide assemblies. Even using existing probes, our imaging strategy should be readily adaptable to helices of different radii, periodicity, peptide sequence, and chirality. Fundamentally, SMOLM enables high-fidelity structural characterization via precise measurements of molecular orientation, rather than via ångström-level imaging resolution, but we note that SMOLM orientation data ideally complement traditional spatial imaging. For example, if two targets of interest produce nearly identical orientation distributions (e.g., KFE8^L^ and KFE8^D^ in Fig. 2) and similar surface hydrophobicity^27^, then sub-nanometer imaging resolution^71,72^ will be critical for resolving structural variations.

Our demonstration of SMOLM for characterizing the nanoscale structure of β-sheet assemblies paves the way for future experimental structure-function studies of self-assembling peptides in native conditions without mutating or permanently modifying the peptides themselves. We remark that the origin and implications of the sinusoidal variation in NR wobble along fibrils of KFE8 and Aβ42 (Fig. 2b iv-vi) are still open questions. Further experimental characterization and theoretical studies of how fluorophore wobble is related to supramolecular structure will be helpful. We look forward to further SMOLM development to enable detailed investigation of polymorphic structures and their impact on the design and function of next-generation biomaterials^73,74^.

## Materials and methods

### Sample preparation and imaging using polarized vortex SMOLM

KFE8^L^ (Ac-FKFEFKFE-NH2) and KFE8^D^ (Ac-fkfefkfe-NH2) peptides were synthesized using standard 9-fluorenylmethoxycarbonyl (Fmoc) chemistry and purified as detailed in our previous publication^34^. To prepare a KFE8 stock solution for TAB imaging, we dissolved lyophilized KFE8 directly into ultrapure water to a concentration of 0.2 mM at room temperature. Aβ42 peptide (DAEFRHDSGYEVHHQKLVFFAEDVGS NKGAIIGLMVGGVVIA) was synthesized and purified by Dr. James I. Elliott (ERI Amyloid Laboratory, Oxford, CT). An Aβ42 stock solution was prepared for imaging as described previously^22,23^. Briefly, we suspended the purified amyloid powder in hexafluoro-2-propanol (HFIP) and sonicated at room temperature for 1 h. Then, the solution was lyophilized. Next, we further purify he amyloid precursors by dissolving the lyophilized Aβ42 in 10 mM NaOH, followed by a 25 min cold water bath. The solution was filtered through a 0.22 μm and a 30 kDa centrifugal membrane filter (Millipore Sigma, UFC30GV and UFC5030). The amyloid fibrils were prepared by incubating 10 μM monomeric Aβ42 in phosphate-buffered saline (PBS, 150 mM NaCl, 50 mM Na_3_PO_4_, pH 7.4) at 37 °C with 200 rpm agitation for 24 h. We found the stock KFE8 and Aβ42 solutions to be stable at 4°C over one week, similar to previous reports^13,75^. All imaging experiments were completed within one week of preparing these stock solutions.

For SMOLM experiments, the KFE8 solution was further diluted to 2 μ M with ultrapure water to observe isolated fibrils. We diluted A β 42 to a concentration of 10 μM in ultrapure water for imaging individual fibrils. For imaging, 10 μL of the solution containing β-sheet assemblies was added to ozone-cleaned 8-well chambered cover glass (high-tolerance No. 1.5, Cellvis, C8-1.5H-N) and let stand for 1 h to allow fibrils to adsorb to the glass. Afterward, we rinsed gently using ultrapure water for less than 10 s before drying. Next, 200 μ L of a solution containing 25 nM Nile red (Fisher Scientific, AC415711000) in ultrapure water was added gently for imaging.

Imaging experiments were performed using our custom-built epifluorescence microscope equipped with the polarized vortex dipole-spread function (DSF)^23^. A circularly polarized 532-nm laser (∼0.73 kW/cm^2^ peak intensity) was used to illuminate the sample at normal incidence. A 1.4 NA oil-immersion objective lens (Olympus UPlan-SApo 100×), dichroic beamsplitter (Chroma, ZT532/640rpc-UF1), and bandpass filter (Semrock, FF01-593/46) were used to collect and isolate fluorescence from NR within the emission path of the microscope. Polarization-sensitive fluorescence detection using the vortex DSF was implemented using a 4f optical system, polarizing beamsplitter (PBS), and liquid-crystal spatial light modulator (SLM, Meadowlark, HSPDM256-520-700-P8) as previously described^23^. Briefly, after the first 4f lens, a PBS splits fluorescence into two orthogonally polarized light paths, and the SLM is placed at the coaligned conjugate back focal plane of both polarization channels. The vortex phase mask is loaded onto the SLM to modulate the NR fluorescence. Finally, the second 4f lens in each channel creates nonoverlapping images of the two polarization channels on a single sCMOS camera (Teledyne photometrics, Prime BSI). For each field of view (FOV), we captured at least 5000 frames with 20 ms exposure time.

### Polarized vortex SMOLM image analysis

The raw images collected by the detector were first converted from AD counts to detected photons by subtracting the camera offset and multiplying by the camera’s conversion gain. Second, the fluorescence background within the raw images was estimated as exactly described by^22^. Briefly, the raw image stacks were separated into 200-frame substacks, and one averaged image was calculated from each substack. Each row and column of the averaged images were fitted to the sum of two 1D Gaussian functions, which in total contain six parameters, including two amplitudes, two centers, and two standard deviations. The averages of the row-wise and column-wise Gaussian fits were used for spatial filtering, which was realized by a biorthogonal wavelet filter (*wavedec2* and *wdencmp* functions in Matlab with parameter level 6 of ‘bior6.8’).

Third, we performed image registration between the two orthogonally polarized channels of our microscope. To determine the geometric relationship between two channels, we captured images of sparse fluorescent beads (Thermo Fisher Scientific, FluoSphere, 540/560, F8800) that are spin-coated on an ozone-cleaned coverglass. The localizations of the beads were determined using the ThunderSTORM plugin^76^ within ImageJ^77^. Localizations of the same bead from multiple consecutive frames were averaged to improve the localization precision. Corresponding control point pairs of each bead across the two channels were used to determine the registration map, which is comprised of the coefficients of a 2D polynomial transformation function (*fitgeotrans* function in Matlab). Later, the localizations of single NR molecules with high precision (< 20 nm) in each channel were also used as control points to further refine the registration map for each FOV.

Fourth, to compensate for optical aberrations, we used images of bright fluorescent beads and a maximum-likelihood estimator to retrieve the pupil phase function, as reported in^23,78^. In short, optical aberrations were modeled as a phase mask at the pupil consisting of the sum of Zernike modes. We considered the first 35 Zernike modes, neglecting piston (flat), tilt (vertical tilt), and tip (horizontal tilt). We calibrated the aberrations of the two polarized channels independently.

Fifth, the offset-subtracted images, registration map, and Zernike coefficients were used as inputs to our bespoke sparsity-promoting maximum-likelihood estimator, RoSE-O^45,46^, using the same analysis protocol in our previous work^23^. The direct output of the estimator is a list of localizations, which includes the 2D location (*x, y*), orientational second moment ***m*** = (*m*_*xx*_, *m*_*yy*_, *m*_*zz*_, *m*_*xy*_, *m*_*xz*_, *m*_*yz*_), and signal *s* of each SM. The second moments were further converted to the mean orientation ***μ*** and wobble angle *α* by minimizing (*fmincon* function in Matlab) the weighted square error between the measured second moments ***m*** from RoSE-O and the second moments computed from the current estimate of ***μ*** and *α* . The weighting matrix is a 6 x 6 Fisher information matrix^79^ calculated from the basis images of the polarized vortex SMOLM. The raw fluorescence images (Fig. S3, Top) from the polarized vortex SMOLM match well with images (Fig. S3, Middle) reconstructed from the measured orientations (Fig. S3, Bottom), demonstrating the high performance of the polarized vortex SMOLM and the bespoke estimator.

When analyzing SMOLM images, the long axis of each fibril was oriented randomly relative to laboratory coordinates. For each region of interest, the long axis of the fibrils was measured by fitting a collection of (*x, y*) positions, corresponding to NR blinking along each fibril, using a linear model (*fit* function in Matlab). To ensure accuracy and precision, we applied a signal filter *s* ≥ 300 photons before linear fitting. The measured long axis for each fibril was then used to generate rotation matrices; after planar rotation transformation, the fibril’s long axis is aligned along the *x* direction, and the NR orientations pointing parallel to the fibril’s long axis are parallel to ***u*** = (1,0,0) . For clarity and convenience, we refer to the newly aligned orientation coordinate system using ***u*** = (*u*_*x*_, *u*_*y*_, *u*_*z*_).

### Theoretical model linking the chirality and periodicity of supramolecular helices to SMOLM orientation measurements

As shown in Fig. 1a, i, a helical ribbon is composed of two helices of identical pitch. Given a left-handed (LH) helix aligned along the *x* axis, we may describe it using the parametric equations

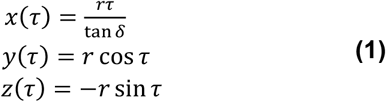

where (*x, y, z*) describes the helical structure in a local coordinate system, *τ* is the helix phase, *r* is the radius of the helix, and *δ* ∈ [0°,90°] is the backbone tilt angle. The parameters *r* and *δ* are determined by the geometry of each helix.

Given that NR binds parallel to the backbone of the helix, the orientations of blinking SMs measured by SMOLM are tangent to the helix and are given by

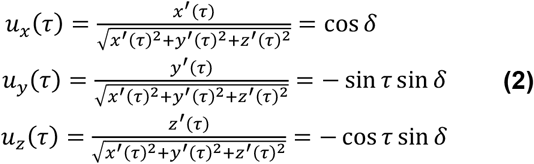

where (*u*_*x*_, *u*_*y*_, *u*_*z*_) represent orientation coordinates with ∥*u*_*x*_, *u*_*y*_, *u*_*z*_ ∥= 1 (Fig. 1a, ii). Notice that *u*_*x*_ is parallel to the long axis of the fibril.

Similarly, a right-handed (RH) helix aligned along the *x* axis and the associated orientations of dyes transiently binding to it can be described by

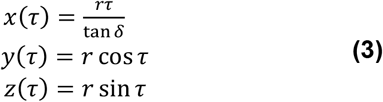

and

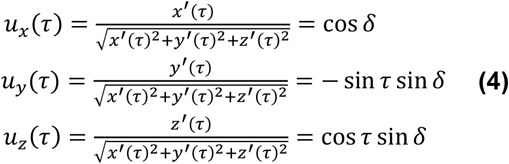

From the above equations, we find that the orientation distributions of LH and RH helices look the same if we neglect the connection between *τ* and the spatial coordinates. That is, accumulating the orientations of all bound SMs along at least one helical period corresponds to a single circle in the orientation domain (Fig. 1a ii and iv).

Interestingly, given sufficient 3D localization precision relative to the radius *r* and period of a helix, SMOLM has the potential to reveal the chirality and the periodicity of a supramolecular helix. For example, when *τ* ∈ [*π*, 2*π*] for an LH helix (Fig. 1a, i), the structure is at its largest height above the coverslip, and the NR orientations from this region lie on the positive *u*_*y*_ ≥ 0 semicircle in the *u*_*x*_ ≥ 0 orientation hemisphere (Fig. 1a, ii). The opposite trend holds for RH helices (Fig 1a, iii and iv). Thus, theoretically, if the height information can be measured with high accuracy and precision, the connection between helix phase *τ*, helix handedness, and NR orientation distribution can be exploited.

Similarly, the periodicity of the helical structure can be extracted from analyzing the 3D positions of NR binding to each helix, or equivalently, the projection of the helix to a sinusoid in 2D given by (*x*(*τ*), *y*(*τ*)) or ( *x*(*τ*), *z*(*τ*) ) in Eqns. 1 and 3. For simplicity, the parameter *τ* is omitted in the following statements. By comparing the parametric equations (Eqs. 1 and 2 or Eqs. 3 and 4) across the spatial and orientation domains, we find that the helix period is also encoded in position-orientation pairs (*x, u*_*y*_) and (*x, u*_*z*_), with identical period to the position-only pairs (*x, y*) and (*x, z*). In addition to these pairs, one may combine all position-orientation measurements for better signal-to-noise ratio and more precise periodicity measurement.

Both chirality and periodicity are currently hidden in SMOLM data by relatively dim NR blinking events, which cause poor localization precision (∼ 12 nm in *xy*)^23^. Future work can improve localization precision by designing more advanced adaptive imaging systems and utilizing brighter dyes.

### Helical optimization algorithm

To determine the helical structure of the β -sheet assemblies (Fig. S4), first, we collect the orientation and rotational mobility measurements of each fluorescent probe along each fibril. The locations are aligned into a common coordinate system, with the fibril’s long axis oriented along the *x* axis and *u*_*x*_ direction, as described above for the convenience of analysis and interpretation.

Second, we measure structure information by fitting the parameter(s) of a specific helix model to a measured orientation distribution (Fig. S4a, iii and iv). This step is formulated as an optimization problem. For a helical bilayer, we solve

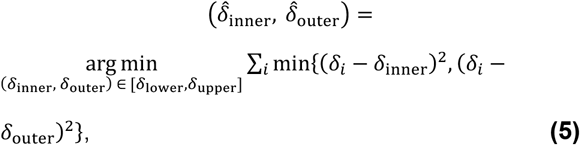

where 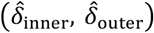 are the resulting estimates of the inner and outer backbone tilt angles, respectively, *δ*_i_ represents the *i*^th^ measurement of the backbone tilt angle from a single NR localization, and *i* is an index over all localizations on a particular fibril. We calculate *δ*_i_ from ***u***_i_ using *δ* = cos^-1^ *u*_*x*_ . If we assume that there is only one layer, i.e., a helical monolayer, we solve

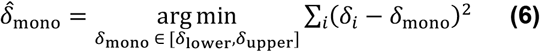

To ensure physically realistic measurements, we constrained the optimization problem using *δ*_lower_ and *δ*_upper_ extracted from our SMOLM data. We use morphological erosion (*medfilt2* function in Matlab) to filter out isolated data points with either small or large *δ* (Fig. S4b and c). Then, we compute values of *δ*_lower_ and *δ*_upper_ such that the annulus in orientation space bounded by *δ*_lower_ and *δ*_upper_ contains all remaining tilt angle measurements *δ* . For KFE8, we set the optimization constraints *δ*_lower_ = 29.23° and *δ*_upper_ = 68.09°. For Aβ42, we set *δ*_lower_ = 0° and *δ*_upper_ = 21.23°.

For structure quantification using experimental data, we used a compound filter, i.e., signal *s* ≥ 500 photons and wobble cone half angle *α*≤ 30°. The signal filter ensures the accuracy and precision of the tilt angle measurements, and we use the wobble cone filter to ensure that the geometry of the helical structure is inferred from NR molecules that are not wobbling excessively during a camera frame. Notably, SMs with large wobble freedom still sense the chemical environment near each helix. Thus, these data are preserved for wobble cone analysis (e.g., Fig. 2b), which quantifies the NR binding behaviors near each fibril.

## Data availability

Raw data and simulation data are available via OSF (https://osf.io/9mf2z/?view_only=3e95ef16e90046dda63e55ade3d5fe93) and from the authors upon reasonable request.

## Code availability

Analysis software are available via OSF (https://osf.io/9mf2z/?view_only=3e95ef16e90046dda63e55ade3d5fe93) and from the authors upon reasonable request.

## Acknowledgements

The authors thank Tingting Wu for inspiring discussions and helpful comments. This work was supported by the National Science Foundation under award number 2047517 to J. S. R. and by the National Institute of General Medical Sciences of the National Institutes of Health under grant number R35GM124858 to M. D. L.

## Author Contributions

W. Z. performed research and analyzed data; W. Z. and C. L. O. prepared samples; T. D. and O. Z. contributed imaging and analytic tools; W. Z., C. L. O., J. S. R., and M. D. L. designed research; J. S. R. and M. D. L. supervised research; W. Z., J. S. R., and M. D. L. wrote the paper with input from all authors.

## Competing Interests

The authors declare no competing interest.

## Notes

https://osf.io/9mf2z/?view_only=3e95ef16e90046dda63e55ade3d5fe93

### Competing Interest Statement

The authors have declared no competing interest.

