## Supporting Information for "Resolving the nanoscale structure of β-sheet assemblies using single-molecule orientation-localization microscopy"

**This PDF file includes:**

Figures S1 to S13
Tables S1

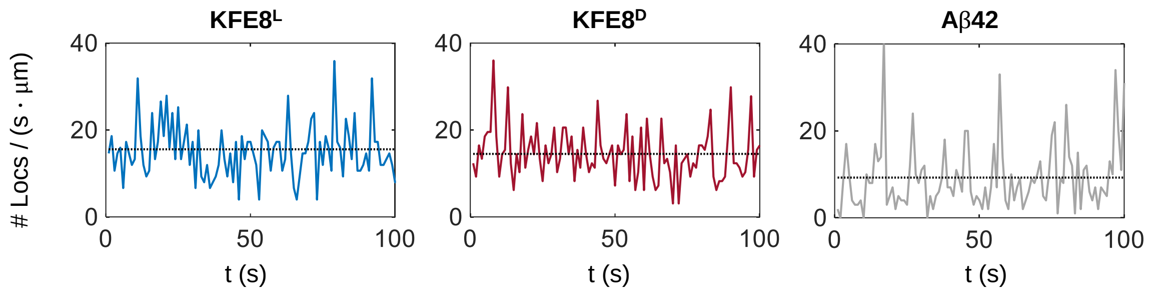


**Fig. S1**. Number of Nile red (NR) localizations over time from the fields of view (FOVs) shown in Fig. 1a-d. (Left) KFE8$^{L}$. (Middle) KFE8$^{D}$. (Right) Aβ42. The localization rate is normalized to the length of each β-sheet fibril. Dotted lines are the mean values of localizations / (second ·µm).

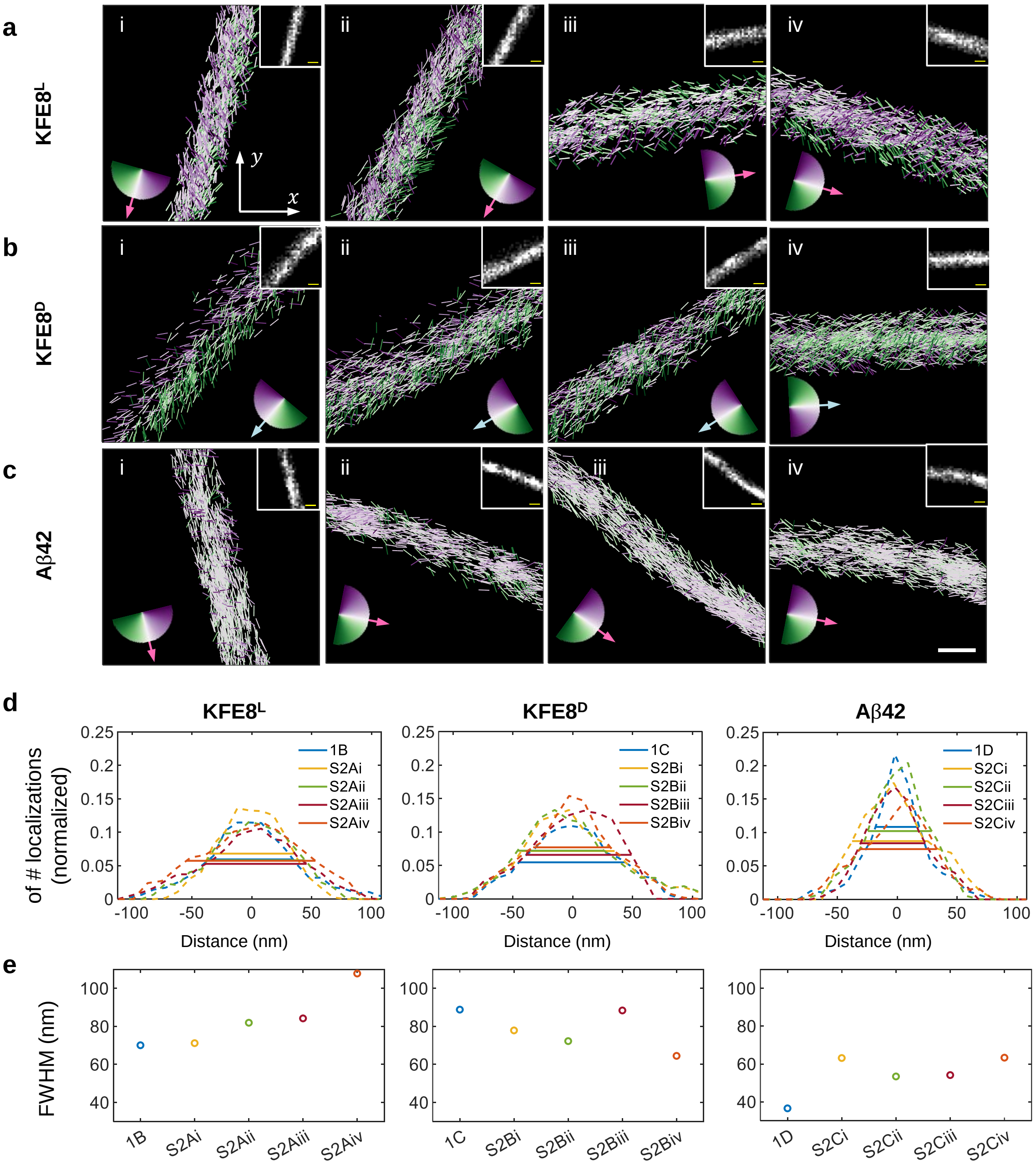


**Fig. S2. a-c,** SMOLM images of (**a**) KFE8$^L$, (**b**) KFE8$^D$, and (**c**) Aβ42. Four representative fibrils are shown in (i-iv) for each peptide. Each NR localization is depicted as a line segment whose color and orientation are set by the projection of the molecule's 3D orientation into the $xy$ plane relative to the long axis $\boldsymbol{u}_x$ of the fibril (pink arrow in **a** and **c**, blue arrow in **b**). Insets, superresolution SMLM images. Scale bars are 100 nm. **d,** Cross-sections of each fibril measured from the SMLM images in **a-c** and Fig. 1b-d. **e,** Full-width at half-maximum (FWHM) fibril thicknesses from **d**.

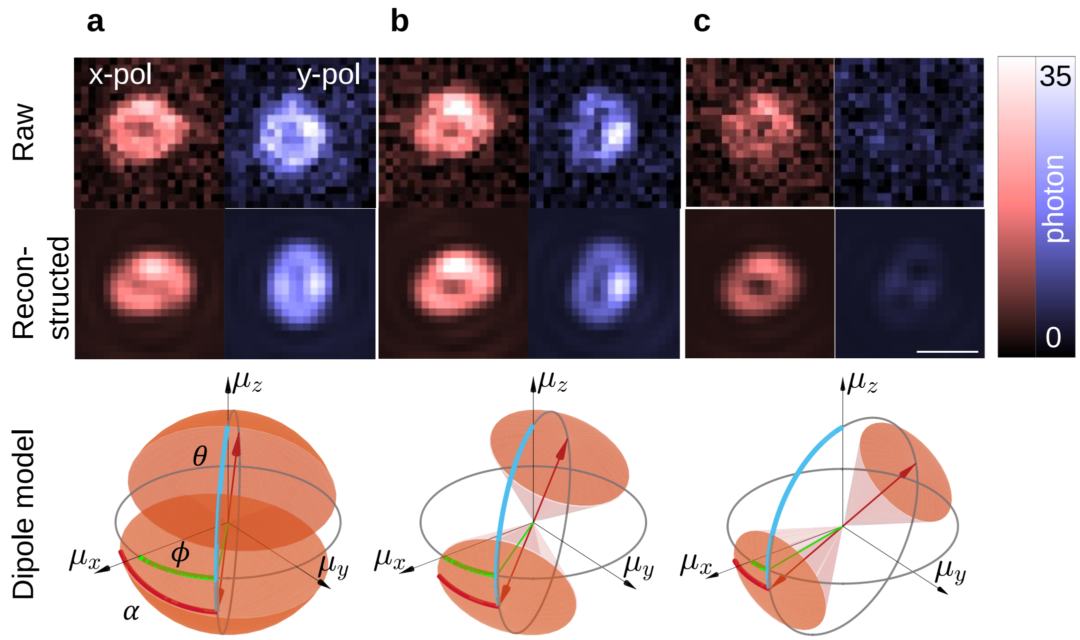


**Fig. S3. a-c**, (Top) Experimental raw (*red*) x-polarized and (*blue*) y-polarized images of a single NR molecule from the polarized vortex microscope[1]. (Middle) Images reconstructed from RoSE-O[2] analysis of the raw images and (Bottom) corresponding visualizations of the NR dipole's orientation and rotational diffusion based on RoSE-O analysis. In each dipole model, the orange cone is the region within which NR "wobbled" during the camera's integration time. The blue and green arcs represent the polar angle and azimuthal angles, respectively. $\mu_x = \sin\theta\cos\phi$, $\mu_y = \sin\theta\sin\phi$, $\mu_z = \cos\theta$. (**a**) $[\mu_x, \mu_y, \mu_z]$ = [0.63, 0.67, -0.40], $\alpha$ = 68°. (**b**) $[\mu_x, \mu_y, \mu_z]$ = [0.78, 0.52, -0.35], $\alpha$ = 41°. (**c**) $[\mu_x, \mu_y, \mu_z]$ = [0.96, 0.21, -0.20], $\alpha$ = 31°. Scale bar is 500 nm.

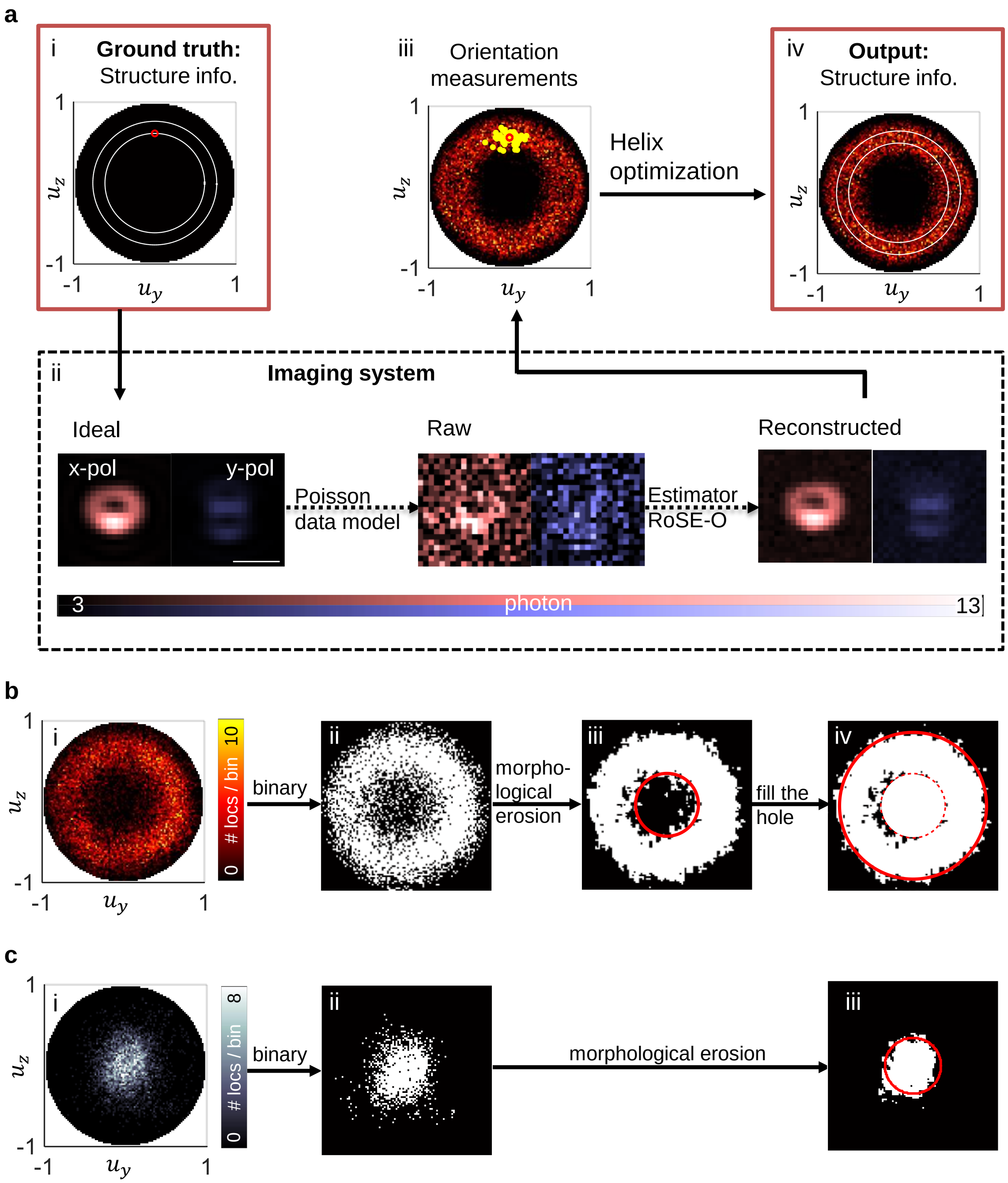


**Fig. S4. a**, Flowchart describing our helix optimization algorithm and its validation. (i) NR orientations are directly linked to the structure of helical bilayers, where $u_x$ is the long axis of the fibril. The two concentric white circles represent the inner $\delta_{\text{inner}}$ and outer $\delta_{\text{outer}}$ backbone tilt angles of the helical bilayer. The red circle is one example binding orientation of NR along the helical structure. (ii) Imaging of a single NR molecule using the polarized vortex dipole-spread function. (*Red*) x-polarized and (*blue*) y-polarized images captured simultaneously by the vortex microscope. (Left) Ideal images of a dipole at the orientation denoted by the red circle in i. Scale bar is 500 nm. (Middle) Simulated noisy raw images with 500 signal photons detected and 3.5 background photons per pixel. (Right) Images reconstructed from RoSE-O$^2$ analysis of the noisy raw images. (iii) Orientation measurements of the helical bilayer in i after passing through the imaging system in ii. The small red circle is the ground-truth orientation in i. The yellow dots correspond to a realistic distribution of noisy measurements matching the signal and background of our experiments. (iv) Orientation measurements overlaid with the estimated backbone tilt angles (white circles) of the helical bilayer using the proposed helix optimization algorithm (See Materials and Methods for more details). The helix optimization algorithm estimates structure information in iv from either simulated or experimental orientation distributions in iii. **b** and **c**, Flowchart describing the determination of constraints in the helix optimization of KFE8 and Aβ42. (i) Accumulated orientation distributions of $KFE8^L$ and $KFE8^D$ (same data as Fig. 2a, Top). (ii) Binarized orientation distribution. (**b**, iii) After morphological erosion, the lower boundary $\delta_{\text{lower}}$ (red circle) bounds the area of the non-zero region from the inside. (**b** iv and **c** iii) The upper boundary $\delta_{\text{upper}}$ (red solid circle) bounds the area of the non-zero region from the outside.

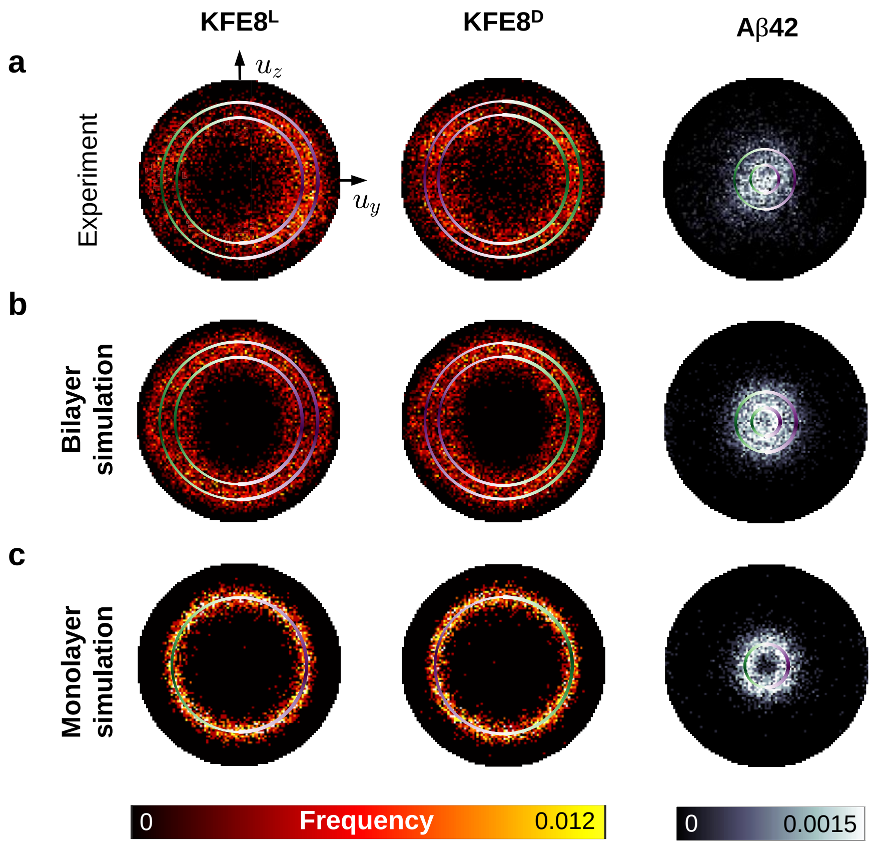


**Fig. S5.** Comparing (**a**) experimental NR orientation distributions from (Left) KFE8$^L$, (Middle) KFE8$^D$, and (Right) Aβ42 to simulated models of (**b**) a helical bilayer and (**c**) a helical monolayer. Plots in **a** and **b** are reproduced from Fig. 2. Ground truth orientation parameters for **b** and **c** are derived from the experimental data in **a.** The circles, i.e., the measured backbone tilt angles, are color-coded according to the chirality of the assemblies (See Fig. 1a). The orientation distributions are binned using $u_y$ = (-1,-0.98,…,1) and $u_z$ = (-1,-0.98,…,1).

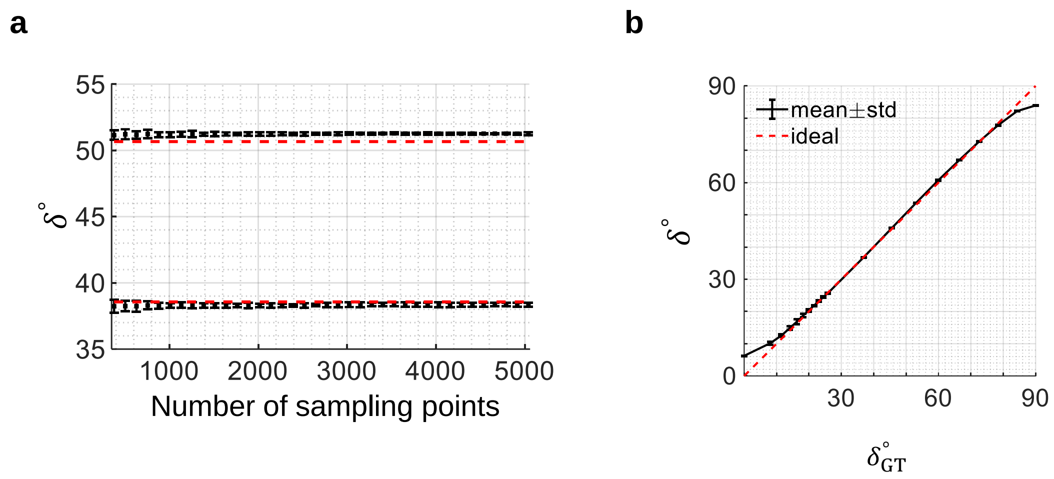


**Fig. S6.** Helix optimization performance. **a**, Accuracy and precision of fitting a helical bilayer versus the number of sampling points based on the geometry of $\mathrm{KFE8}^{\mathrm{L}}$. Red dashed lines: ground truth, same as Fig. 2a Left. Dots and error bars: mean $\pm$ standard deviation of $\delta_{\mathrm{inner}}$ and $\delta_{\mathrm{outer}}$. **b**, Accuracy and precision of fitting a helical monolayer versus the ground truth backbone tilt angle $\delta_{\mathrm{GT}}$. Red dashed line: ground truth. Line and error bars: mean $\pm$ standard deviation of $\delta$.

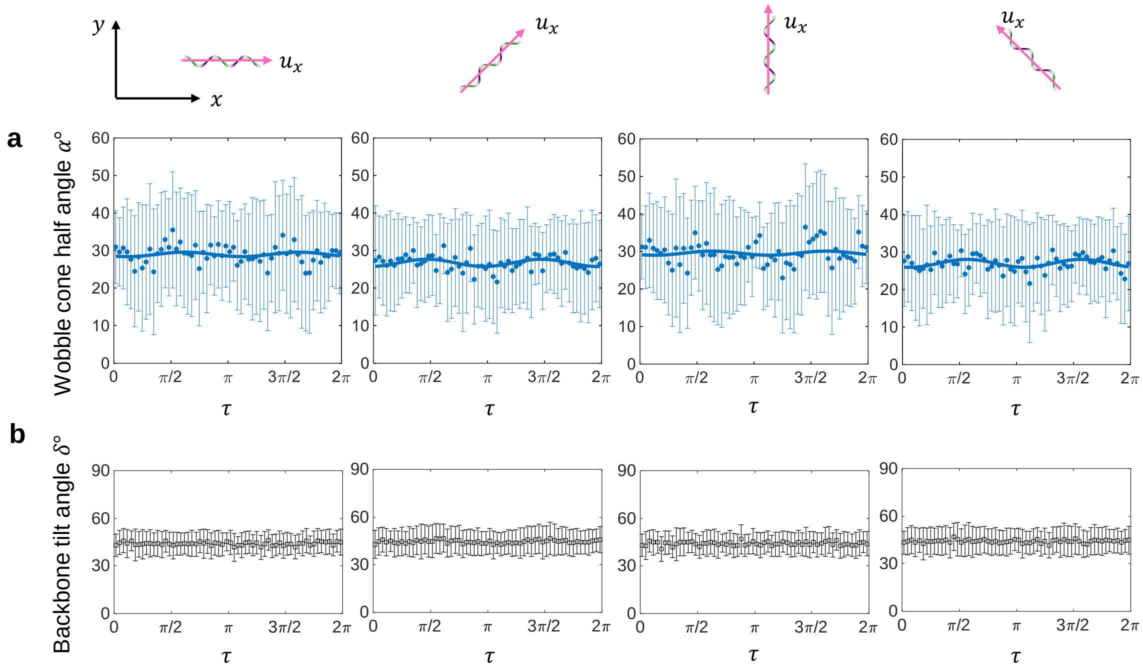


**Fig. S7.** Simulated measurements of helical ribbons whose NR localizations have a constant wobble half-cone angle $\alpha = 30^{\circ}$. Measured (**a**) median wobble angle $\alpha$ and (**b**) median backbone tilt angle $\delta$ distributions with respect to helix phase $\tau$. The simulations were performed according to the flowchart in Fig. S4 and are detailed in Materials and Methods. The simulated fibrils are oriented differently with respect to the $x$ axis of the microscope; from Left to Right, the angles between the long axis $u_x$ of the helical ribbon (pink arrow) and the $x$ axis of the microscope are 0°, 45°, 90°, and 135°, respectively. Black dots and error bars, mean $\pm$ standard deviation. Blue solid lines in **a** are from sinusoidal functions. From Left to Right, the sinusoidal functions are $\alpha = -0.6° \cos(2\tau - 0.78) + 29.0°$, $\alpha = -1.0° \cos(2\tau + 0.18) + 26.7°$, $\alpha = -0.5° \cos(2\tau - 0.83) + 29.5°$, $\alpha = -1.1° \cos(2\tau - 0.42) + 27.0°$. Bin sizes are $\pi/30$, i.e., 6°.

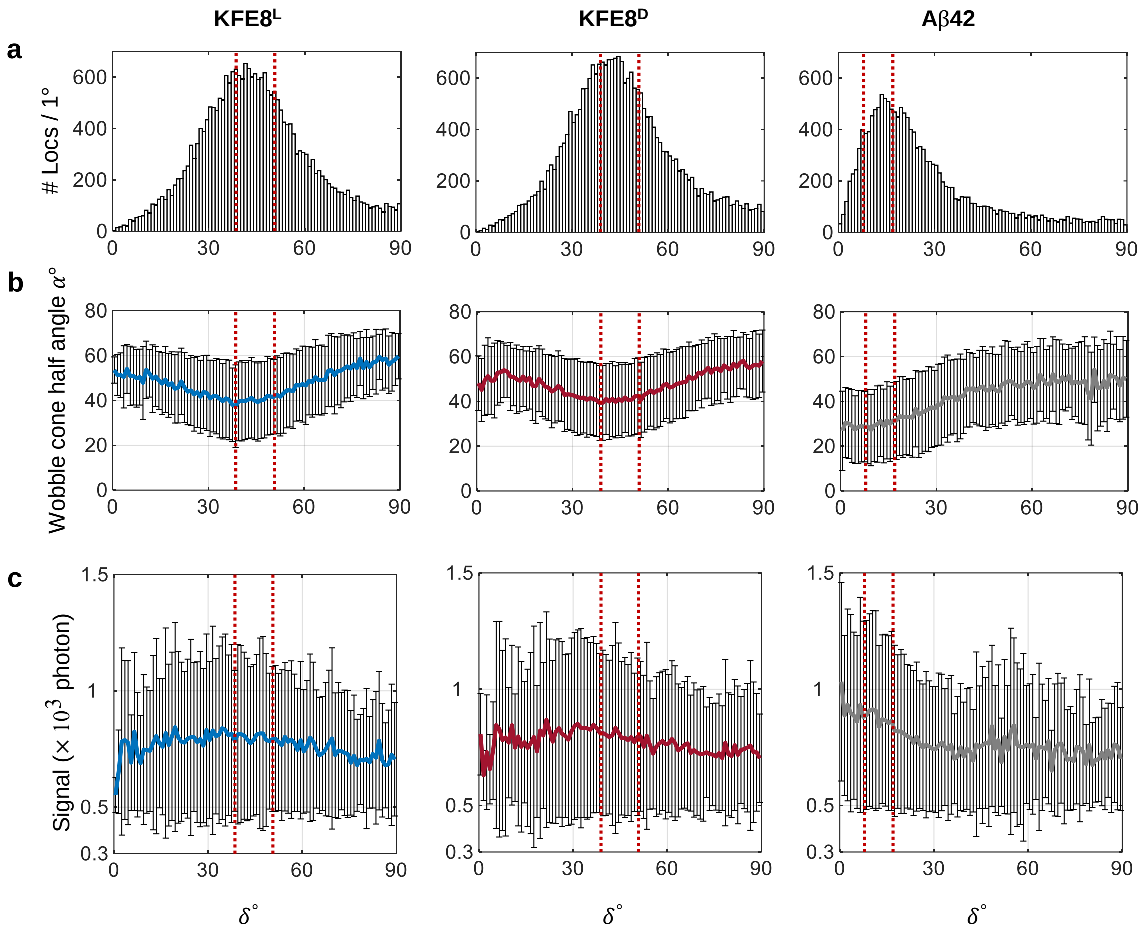


**Fig. S8.** Full distributions of NR (**a**) localization density, (**b**) wobble, and (**c**) detected signal are quantified with respect to backbone tilt angle $\delta$ for (Left) KFE8$^L$, (Middle) KFE8$^D$, and (Right) Aβ42; **a** and **b** match the data in Fig. 2b. Solid lines and error bars, mean $\pm$ standard deviation. Red vertical dotted lines are the backbone tilt angles of the helical ribbons, as estimated by helical optimization, and match the overlaid circles in Fig. 2a. Bin sizes are 1°.

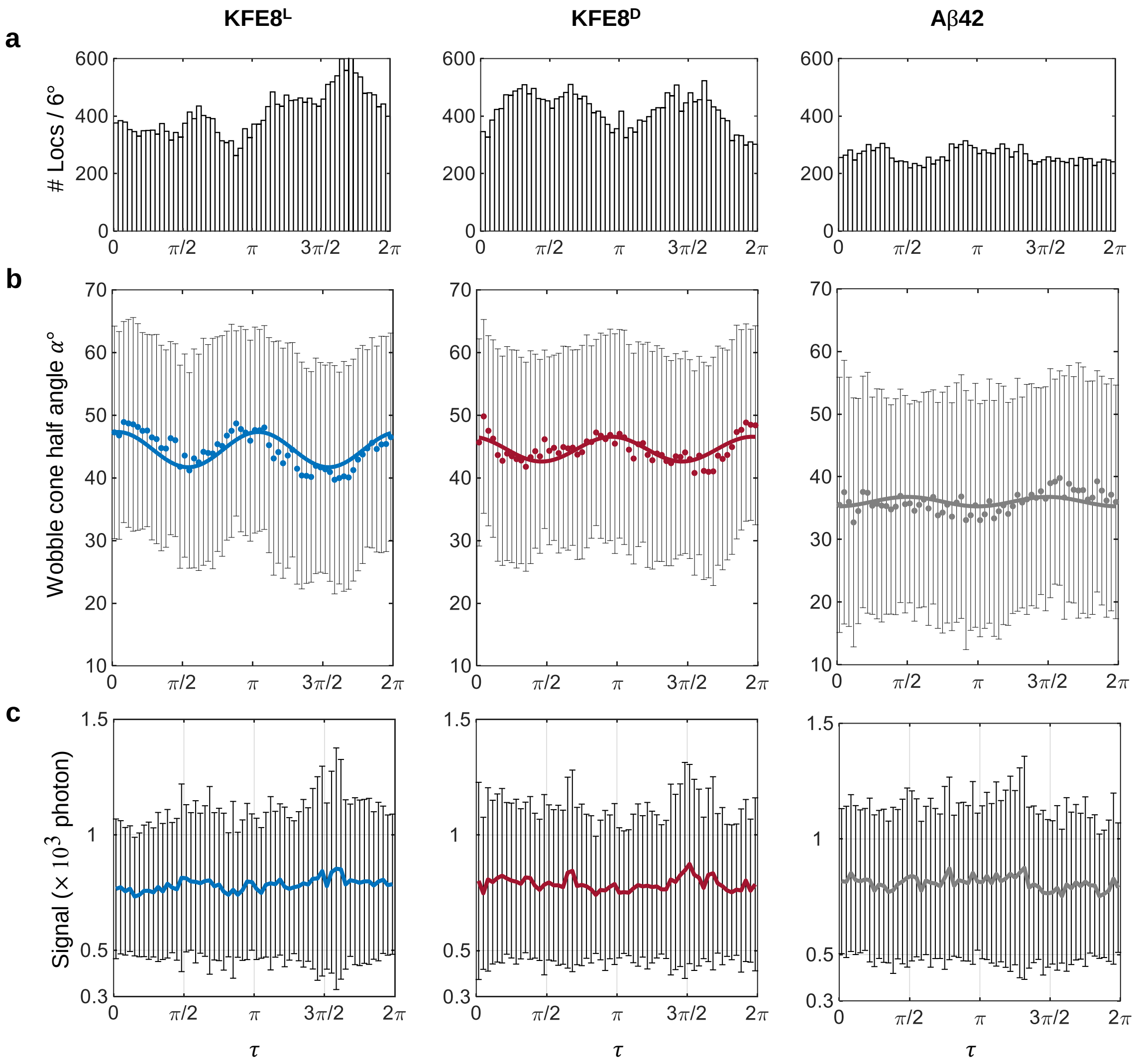


**Fig. S9.** Full distributions of NR (**a**) localization density, (**b**) wobble, and (**c**) detected signal are quantified with respect to helix phase $\tau$ for (Left) KFE8$^{L}$, (Middle) KFE8$^{D}$, and (Right) Aβ42; **a** and **b** match the data in Fig. 2b. Solid lines and error bars, mean $\pm$ standard deviation. Bin sizes are $\pi/30$, i.e., 6°.

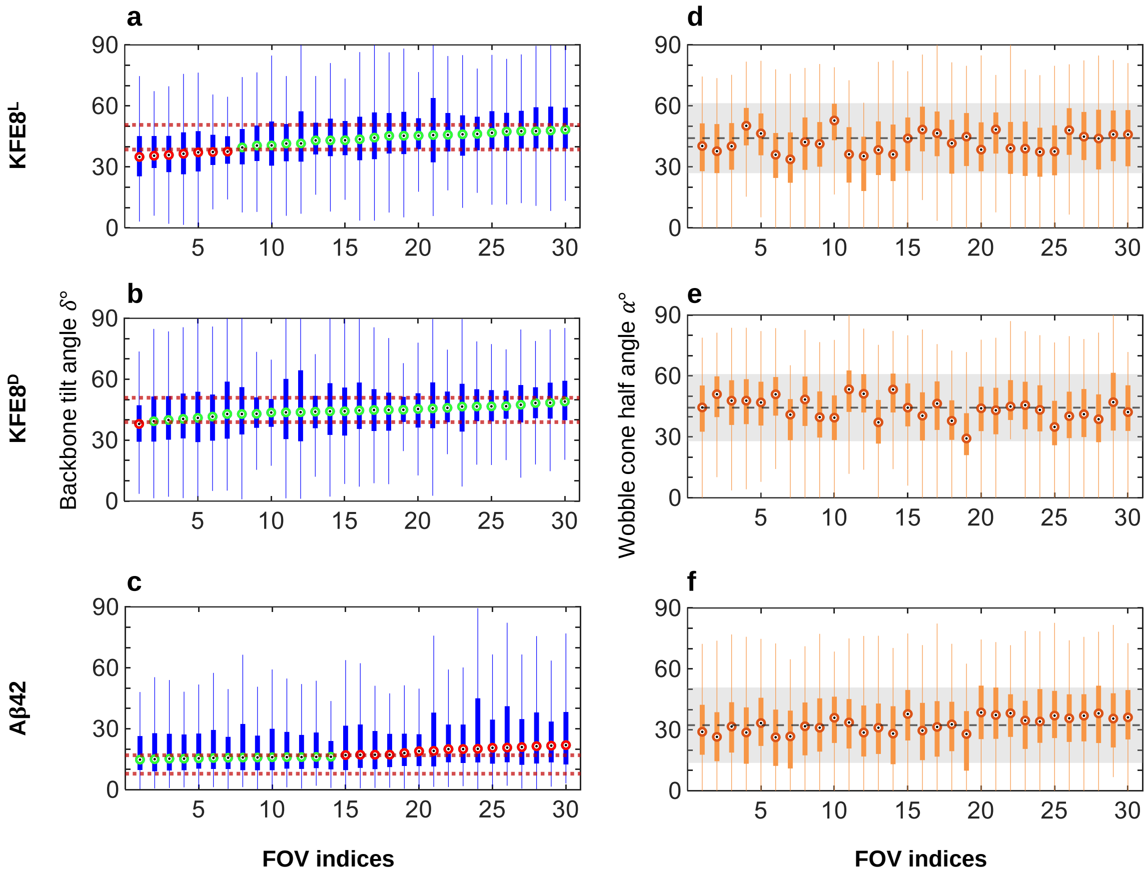


**Fig. S10. a-c**, Full distributions of measured backbone tilt angles $\delta$ for (**a**) KFE8$^L$, (**b**) KFE8$^D$, and (**c**) Aβ42 shown in Fig. 3a-c. Hollow circles are median backbone tilt angles of each FOV. Red dotted lines are the backbone tilt angles of the helical bilayer, as estimated by helical optimization, and match the overlaid concentric circles in Fig. 2a. Green circles lie within the estimated backbone tilt angles of the bilayer; red circles outside the range. **d-f**, Full distributions of measured wobble angles $\alpha$ for (**d**) KFE8$^L$, (**e**) KFE8$^D$, and (**f**) Aβ42 shown in Fig. 3d-f. Orange circles are the median half-cone angles. Gray dashed line and shaded region are the mean $\pm$ standard deviation of the accumulated experimental data in Fig. 2b. Filled boxes: 25$^{th}$ to 75$^{th}$ percentiles. Whiskers extend to the minimum and maximum measured values.

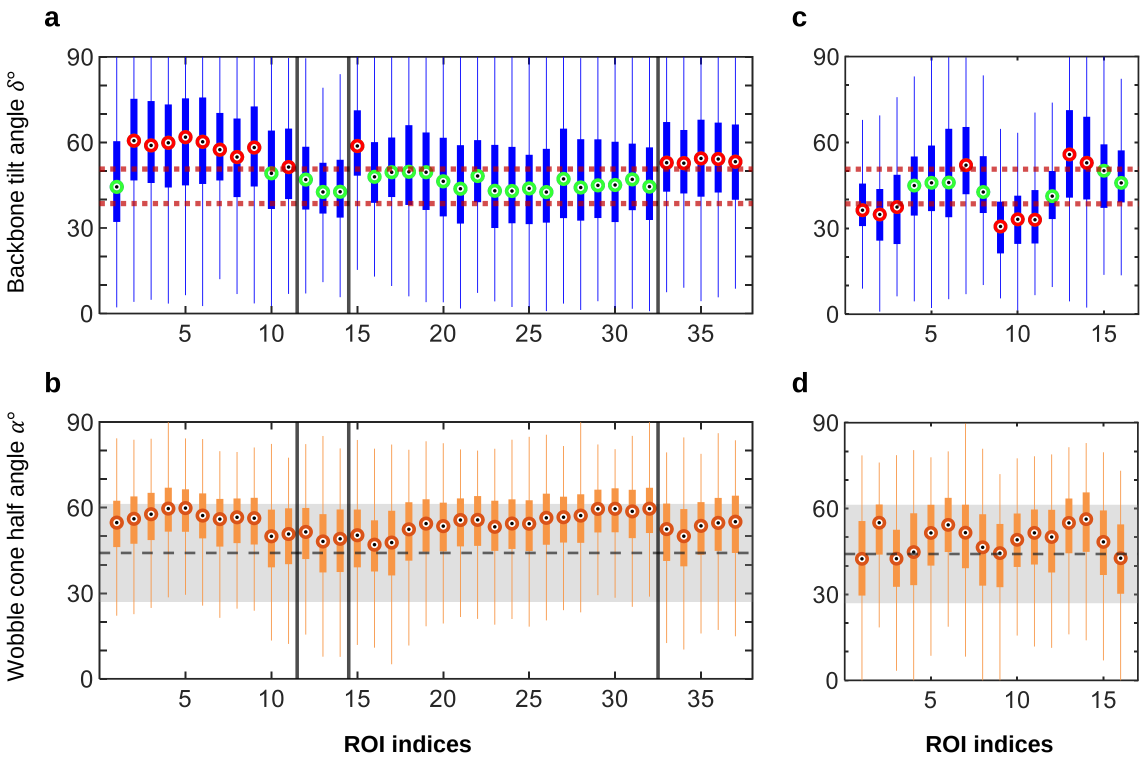


**Fig. S11.** (**a** and **c**) Full distributions of measured backbone tilt angles $\delta$ in Fig. 4b and e. Hollow circles are median backbone tilt angles of each FOV. Dashed lines are the backbone tilt angles of the helical bilayer, as estimated by helical optimization, and match the overlaid concentric circles in Fig. 2a Left. Green circles lie within the estimated backbone tilt angles of the bilayer; red circles outside the range. (**b** and **d**) Full distributions of wobble angles $\alpha$ in Fig. 4c and f. Orange circles are the median wobble cone half angles. Gray dashed line and shaded region are the mean $\pm$ standard deviation of the accumulated experimental data in Fig. 2b. Filled boxes: 25$^{th}$ to 75$^{th}$ percentiles. Whiskers extend to the minimum and maximum measured values.

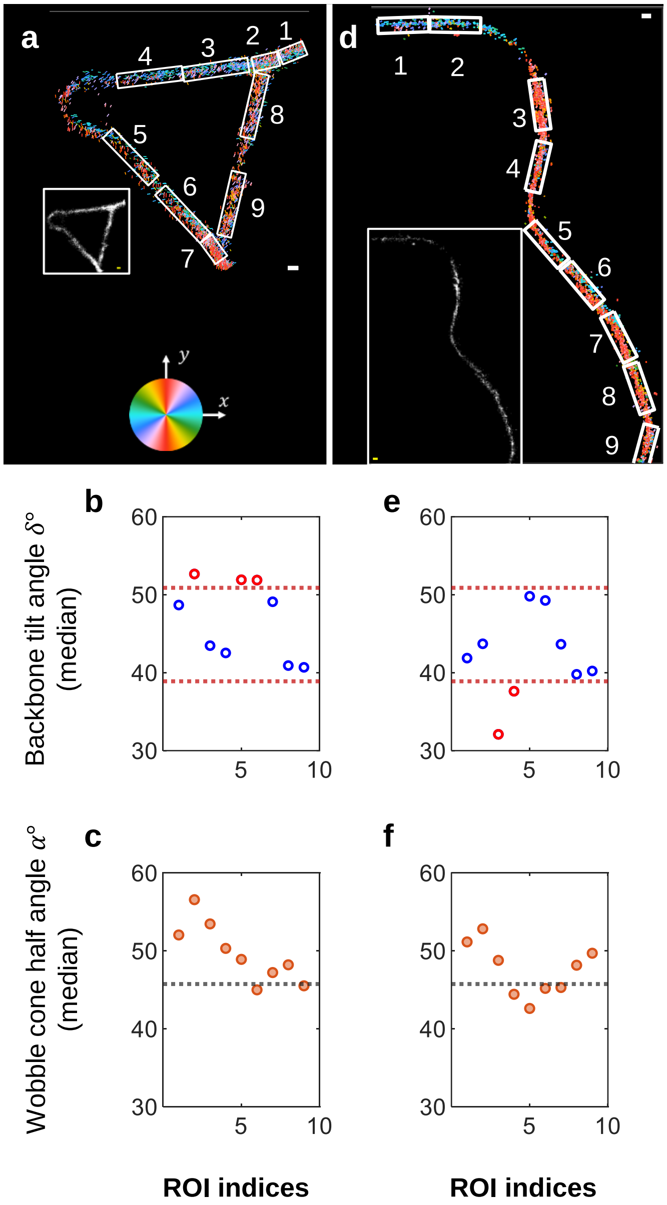


**Fig. S12.** Structural polymorphism of KFE8$^{D}$ revealed by SMOLM: **a-c**, Branching and **d-f**, curved structure structures. (**a** and **d**) SMOLM and (Inset) SMLM images. NR orientations in the SMOLM images are represented as colored lines that are color-coded and oriented according to the projection of the 3D NR orientation into the $xy$ plane relative to the $\mu_x$ axis, which is defined by the microscope instead of the long axis of the fibril. Scale bars are 100 nm. (**b** and **e**) Median backbone tilt angles $\delta$ within each region of interest (ROI), which are shown in SMOLM images. Red dotted lines are the backbone tilt angles of the helical bilayer, as estimated by helical optimization, and match the overlaid concentric circles in Fig. 2a Middle. (**c** and **f**) Median wobble angle $\alpha$ (orange circle) within each ROI. Black dotted lines are the median of the accumulated experimental data in Fig. 2b.

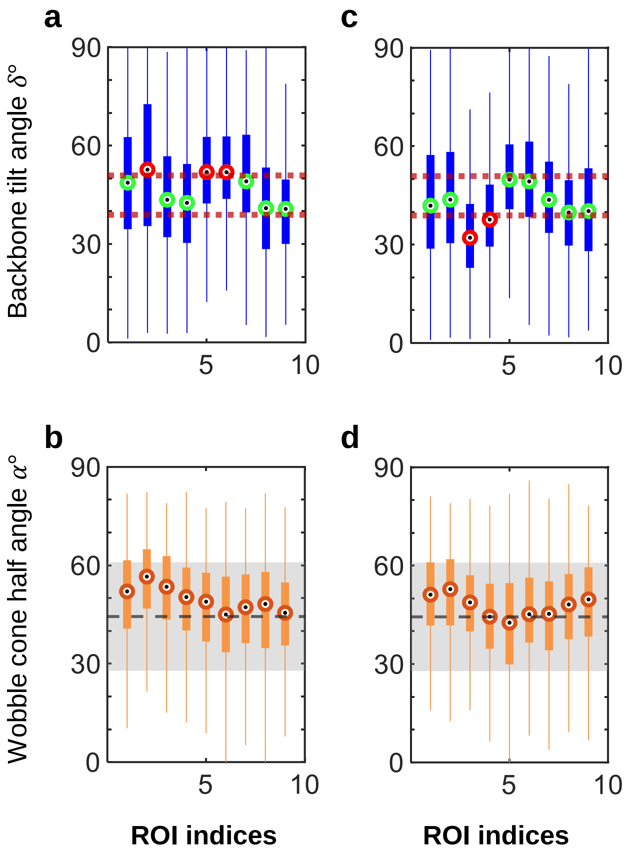


**Fig. S13.** (**a** and **c**) Full distributions of measured backbone tilt angles $\delta$ in Fig. S12b and e. Hollow circles are median backbone tilt angles of each FOV. Dashed lines are the backbone tilt angles of the helical bilayer, as estimated by helical optimization, and match the overlaid concentric circles in Fig. 2a Middle. Green circles lie within the estimated backbone tilt angles of the bilayer; red circles outside the range. (**b** and **d**) Full distributions of wobble angles $\alpha$ in Fig. S12c and f. Orange circles are the median wobble cone half angles. Gray dashed line and shaded region are the mean $\pm$ standard deviation of the accumulated experimental data in Fig. 2b. Filled boxes: 25$^{th}$ to 75$^{th}$ percentiles. Whiskers extend to the minimum and maximum measured values.

**Table S1.** Simulated performance of measuring structural parameters using helical bilayer optimization. (**a**) Accuracy of outer layer estimation, $\delta_{\text{outer}} - \delta_{\text{outer,GT}}$ (°). (**b**) Precision of outer layer estimation, $\sigma_{\delta_{\text{outer}}}$ (°). (**c**) Accuracy of inner layer estimation, $\delta_{\text{inner}} - \delta_{\text{inner,GT}}$ (°). (**d**) Precision of inner layer estimation, $\sigma_{\delta_{\text{inner}}}$ (°). A range of ground-truth outer and inner layer tilt angles, $\delta_{\text{outer}}$ and $\delta_{\text{inner}}$ respectively, were explored. Details are discussed in Materials and Methods.

**a**

| $\delta_{\text{outer,GT}}$ \ $\delta_{\text{inner,GT}}$ | 32 | 34 | 36 | 38 | 40 |
|---|---|---|---|---|---|
| **48** | 0.31 | 0.47 | 0.53 | 0.96 | 1.26 |
| **50** | 0.41 | 0.67 | 0.86 | 1.01 | 1.14 |
| **52** | -0.30 | -0.06 | -0.07 | -0.03 | 0.16 |
| **54** | -0.70 | -0.62 | -0.52 | -0.41 | -0.30 |

**b**

| $\delta_{\text{outer,GT}}$ \ $\delta_{\text{inner,GT}}$ | 32 | 34 | 36 | 38 | 40 |
|---|---|---|---|---|---|
| **48** | 0.81 | 0.82 | 0.93 | 0.88 | 0.89 |
| **50** | 0.77 | 0.73 | 0.84 | 0.84 | 0.94 |
| **52** | 0.47 | 0.54 | 0.77 | 0.69 | 0.73 |
| **54** | 0.53 | 0.61 | 0.55 | 0.54 | 0.62 |

**c**

| $\delta_{\text{outer,GT}}$ \ $\delta_{\text{inner,GT}}$ | **32** | **34** | **36** | **38** | **40** |
|---|---|---|---|---|---|
| 48 | -0.38 | -0.10 | -0.91 | -0.90 | -1.56 |
| 50 | -0.15 | 0.29 | -0.31 | -0.38 | -1.04 |
| 52 | -0.21 | 0.31 | -0.44 | -0.44 | -0.92 |
| 54 | -0.03 | 0.34 | -0.23 | -0.11 | -0.60 |

**d**

| $\delta_{\text{outer,GT}}$ \ $\delta_{\text{inner,GT}}$ | **32** | **34** | **36** | **38** | **40** |
|---|---|---|---|---|---|
| 48 | 0.80 | 0.58 | 0.85 | 0.66 | 1.05 |
| 50 | 0.59 | 0.77 | 0.81 | 0.50 | 0.81 |
| 52 | 0.60 | 0.63 | 1.00 | 0.45 | 0.81 |
| 54 | 0.54 | 0.63 | 0.88 | 0.54 | 0.62 |
